# Microflora Danica: the atlas of Danish environmental microbiomes

**DOI:** 10.1101/2024.06.27.600767

**Authors:** CM Singleton, TBN Jensen, F Delogu, EA Sørensen, VR Jørgensen, SM Karst, Y Yang, KS Knudsen, M Sereika, F Petriglieri, S Knutsson, SM Dall, RH Kirkegaard, JM Kristensen, BJ Woodcroft, DR Speth, STN Aroney, The Microflora Danica Consortium, M Wagner, MKD Dueholm, PH Nielsen, M Albertsen

**Affiliations:** Center for Microbial Communities, Department of Chemistry and Bioscience, Aalborg University, Aalborg, Denmark; Centre for Microbiome Research, School of Biomedical Sciences, Queensland University of Technology (QUT), Translational Research Institute, Woolloongabba, Australia; Centre for Microbiology and Environmental Systems Science, University of Vienna, Vienna, Austria

## Abstract

The last 20 years have witnessed unprecedented advances in revealing the microbiomes underpinning important processes in natural and human associated environments. Recent large-scale metagenome surveys record the variety of microbial life in the oceans^1^, wastewater^2^, human gut^3,4^, and earth^5,6^, with compilations encompassing thousands of public datasets^7–13^. So far, large-scale microbiome studies either miss functional information or consistency in sample processing, and although they may cover thousands of locations, these are missing resolution, sparsely located, or lacking metadata. Here, we present Microflora Danica, an atlas of Danish environmental microbiomes, encompassing 10,686 shotgun metagenomes and 449 full-length 16S and 18S rRNA datasets linked to a detailed 5 level habitat classification scheme. We determine that while human-disturbed habitats have high alpha diversity, the same species reoccur, revealing hidden homogeneity and underlining the importance of natural systems for total species (gamma) diversity. In-depth studies of nitrifiers, a functional group closely linked to climate change, challenge existing perceptions regarding habitat preference and discover several novel nitrifiers as more abundant than canonical nitrifiers. Together, the Microflora Danica dataset provides an unprecedented resource and the foundation for answering fundamental questions underlying microbial ecology: what drives microbial diversity, distribution and function.

## Results

### The Microflora Danica data set

In 1752, King Frederik the 5^th^ of Denmark, known for his “*generous attitude […] towards natural science and applied art*” commissioned the Flora Danica project, initiating an “*Opus Incomparibile*” that took 122 years and produced one of the world’s most unique works in natural history^14^. Over 3000 botanic engravings and 17 volumes were completed of flowers and plants, with which, “*according to the unanimous contention of all connoisseurs*”, “*the whole world can eventually reap all the fruits that follow the extension of a science which, with regard to the benefit of mankind, is one of the most useful and without which medicine and economics would lack important advantages*”^14^. In 2019, we initiated the Microflora Danica (MFD) project with the aim of cataloguing the microbiome of Denmark, in the hopes that the microflora of Denmark can be similarly studied and their riches contribute to the extension of science.

The Microflora Danica data set consists of 10,686 samples that are associated with DNA sequencing, GPS coordinates and a highly curated 5-level MFD Ontology (MFDO) (i.e., habitat classification system; **Figure 1**, **Supplementary Figure 1**) that can be linked to other ontologies (EMPO^5,6^, Natura 2000^15^ and EUNIS^16^). The MFDO1 ontology level represents 24 different categories (**Figure 1c**) and reflects primary Danish land uses: e.g. agricultural fields (3,003 samples), grassland formations (1,393 samples), forests (1,328 samples) and urban greenspaces (711 samples). The unprecedented breadth of sampling is exemplified by our coverage of 27% of the 986 registered lakes in Denmark.

**Figure 1:**
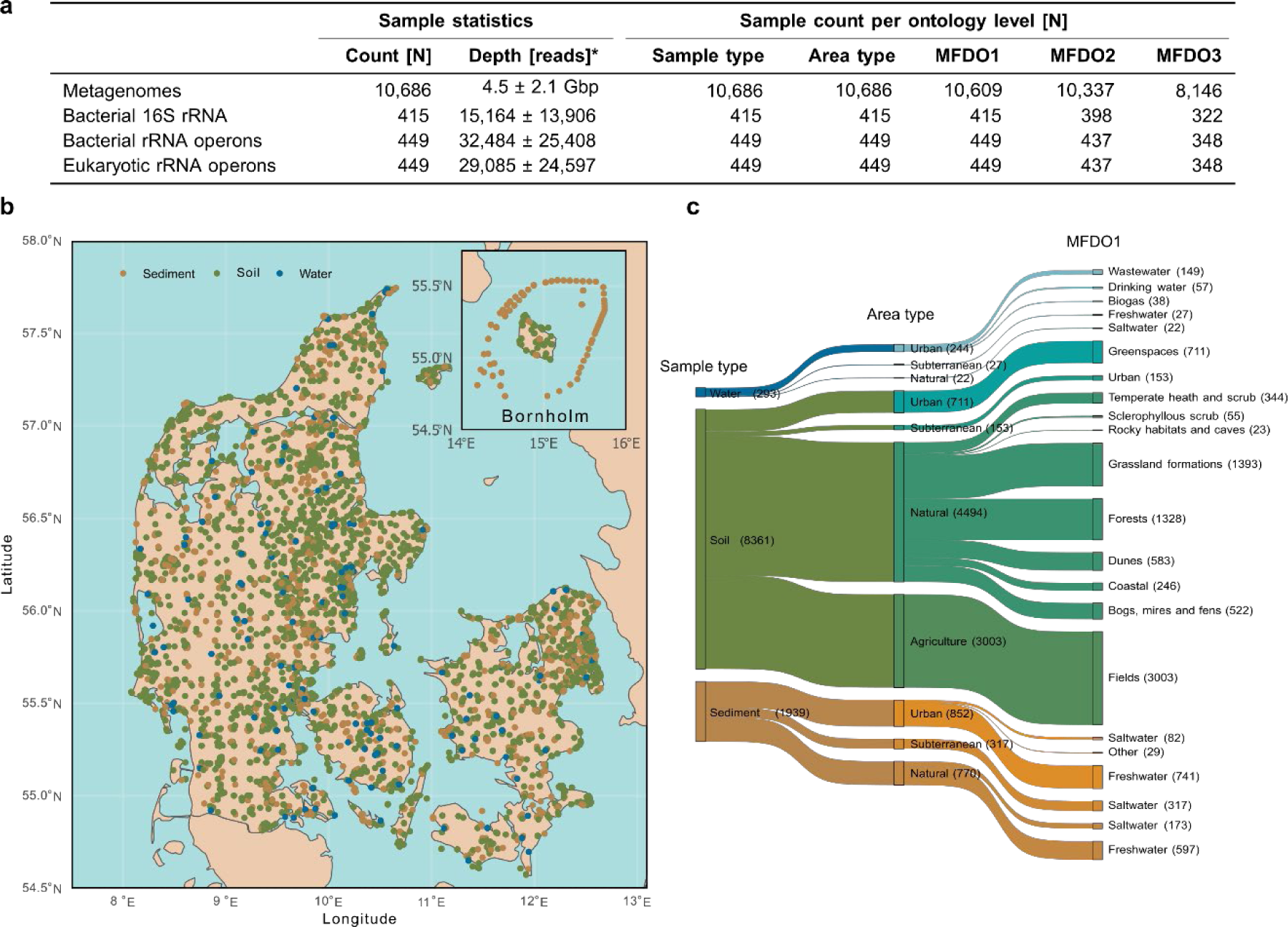
MFD sampling campaign and ontology. **a.** The metagenome and rRNA amplicon data sequencing depths are reported as average ± standard deviation. *The unit of measurement for depth is [reads] except for metagenomes, for which it is reported as Gbp. **b.** The MFD samples cover the land of Denmark and its surrounding waters. The map depicts the locations of the samples used for metagenomics, whilst the colours represent the three different sample types. The upper right cutouts show the island of Bornholm east of the mainland. **c.** Sample count in the first three levels of the habitat ontology. The MFD habitat ontology accounts for a variable number of samples per category/branch. The Sankey reports the first three levels of the ontology and the thickness of the branches is proportional to the number of samples in each category. Only classes with n>20 samples and non-empty MFDO1 classification are reported. Each habitat category is followed by the number of samples for that category in parentheses. **Supplementary Figure 1** shows the Sankey plot including all five levels of the ontology.

To facilitate diversity analysis of bacteria and eukaryotes we selected 449 representative samples for rRNA operon sequencing, generating a total of 14.9 million bacterial (median 4,528 bp) and 13.4 million eukaryotic rRNA operon sequences (median 4,035 bp). Furthermore, we used unique molecular identifiers (UMIs) to sequence 6.4 million near full-length bacterial 16S rRNA genes (median 1,355 bp) from 415 samples, which features an extremely low chimaera rate^17^. This dataset is an order of magnitude larger than the current most comprehensive database SILVA 138.1^18^, which contains 2.1 million 16S rRNA and 0.17 million 18S rRNA gene sequences > 1,200 bp (Bacteria and Eukarya) and 900 bp (Archaea) before clustering. We combined the MFD 16S rRNA gene sequences with the public 16S rRNA gene sequences (SILVA v138.1 SSURef NR99^18^, EMP500^6^, AGP70^17^, MiDAS^2,19^, Askov/Cologne^20^) resulting in a compilation of 30.2 million sequences. These sequences were processed using Autotax^21^, creating the Microflora Global (MFG) database of 350,815 species representatives (clustered at 98.7% nucleotide identity) with a complete 7-level taxonomic string.

Illumina metagenome sequencing of the 10,686 MFD samples generated an average of 4.5 Gbp per sample, totalling 48.2 Tbp of sequencing data (**Figure 1a**). We assembled and binned these metagenomes, creating the metagenome-assembled genomes (MAGs) database that comprises a total of 19,253 metagenome-assembled genomes of at least MIMAG^22^ medium quality, with 5,518 species groups (95% ANI). These MAGs represent a resource for exploring the functional potential of identified core species and the diversity and distribution patterns of microbial guilds (e.g. nitrifying microbes, see below). Combined, the databases, ontology, associated metadata and spatial resolution provide an extraordinary resource to investigate diversity and function questions in microbial ecology.

### The Danish microbiome is diverse and mirrors the global microbiome

We investigated bacterial diversity using the MFD FL16S rRNA dataset of 457 habitat representative samples and 21.3 million sequences (**Figure 2a**, see **Methods**) (**Supplementary Figure 2**) that enable species-level resolution. We observed 168,938 species (98.7% OTUs^23^) across the samples (**Figure 2a,b,c**), and estimated the total richness to be 194,744-196,023 species (95% CI), of which 75,515-75,806 species (95% CI) can be considered common^24,25^. These findings indicate that the MFD FL16S rRNA database captures Denmark’s dominant species in the investigated habitats and sets a minimum estimate of the total free-living bacterial gamma diversity of Denmark. Compared to the SILVA 138.1 database, 75% of the identified MFD OTUs are novel at the species level (**Figure 2b,c**). However, the discovery rate of novelty quickly decreases, indicating that most bacterial orders in the tree-of-life have been discovered at the 16S rRNA gene level (**Figure 2b,c**).

**Figure 2:**
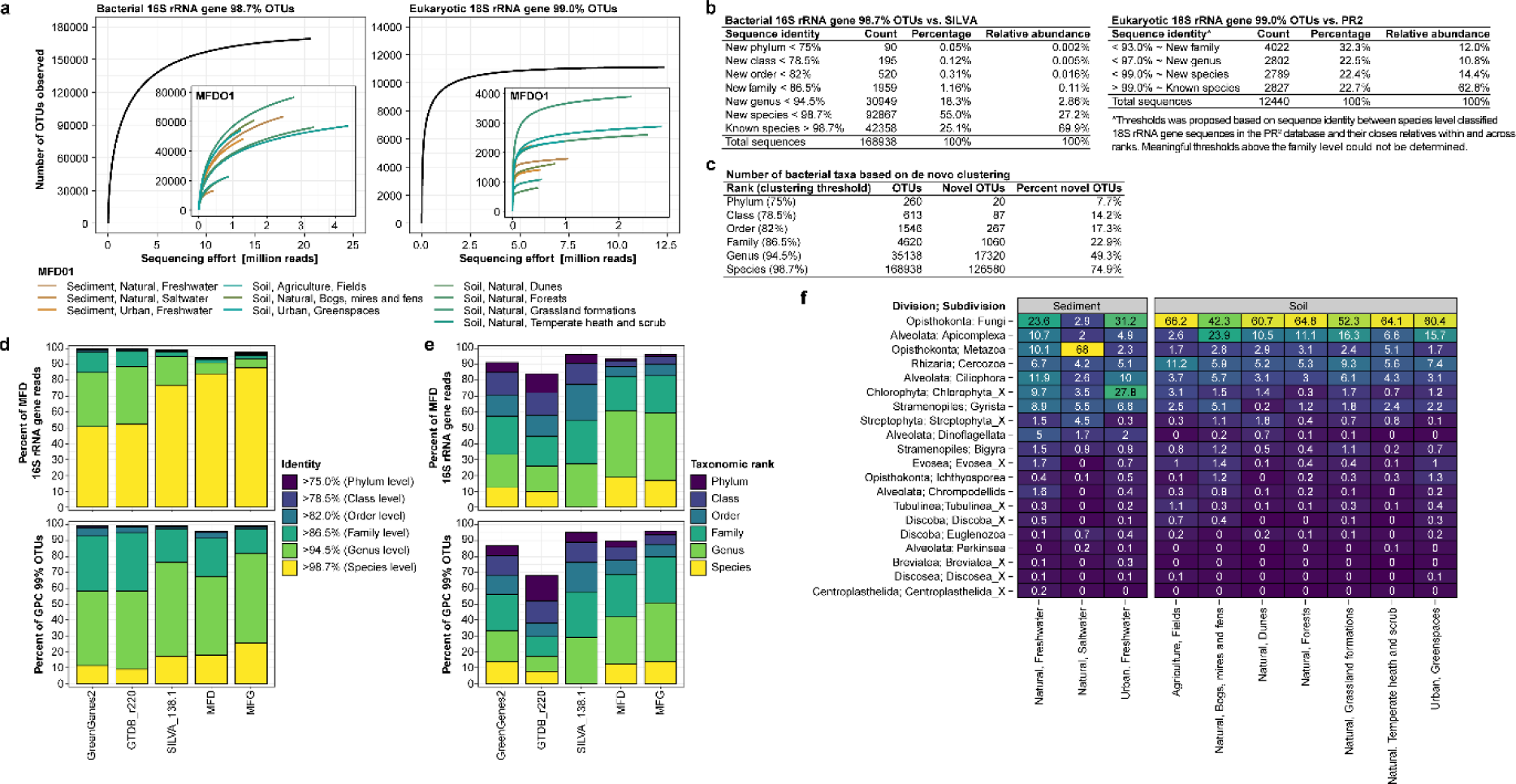
Novelty and diversity based on near full-length 16S and 18S rRNA gene sequencing. **a.** Species-level rarefaction curves of bacterial 16S rRNA and eukaryotic 18S rRNA gene OTUs. **b.** Sequence novelty of species level clustered bacterial 16S rRNA gene OTUs (98.7%) against SILVA v138.1 NR99 and eukaryotic 18S rRNA gene OTUs (99.0%) against PR2^26^ v5.0.0. Taxonomic thresholds for bacteria were adapted from Yarza et al. 2014^23^, whereas those for eukaryotic were calculated using a similar approach based on sequences from the PR^2^ database (**Supplementary Note 1**). **c.** Taxonomic diversity and novelty based on clustering bacterial 16S rRNA genes using the proposed threshold for taxonomic ranks^23^. OTUs were considered novel if the percent identity of the centroid and its closest match in SILVA was below the identity threshold. **d.** Database evaluation based on 16S rRNA gene read extracted from selected MFD metagenomes and V4 OTUs clustered at 99% identity from the Global Prokaryote Census (GPC) dataset^13^. Sequence novelty was determined by non-heuristic mapping of metagenomic reads or OTUs to each reference database. **e.** Classification of metagenomic reads or OTUs was done using the SINTAX classifier^27^. The following databases were used besides the ones created here: GreenGenes2_2022_10 backbone taxonomy^28^, GTDB_ssu_all_r220^29^, SILVA_138.1_SSURef_NR99^18^ **f.** Heatmap displaying the relative abundance of the top 20 subdivisions across MFDO1 habitats based on eukaryotic 18S rRNA gene OTUs.

To evaluate the taxonomic coverage of the MFD and MFG 16S rRNA gene database, we compared the mappings (**Figure 2d**) and classifications (**Figure 2e**) of a representative subset of our 10,686 short-read metagenome-based 16S rRNA gene fragments (see **Methods**) with results using public databases. We were able to classify ∼60% of all extracted 16S rRNA gene reads to genus level using the MFD database, which is ∼80% more than the second best performing database (GreenGenes2) (**Figure 2e**). To determine the global taxonomic coverage of the MFG database, we evaluated it against the Global Prokaryote Census (GPC) dataset^13^. MFG had better coverage at all taxonomic levels compared to the other databases (**Figure 2d**), and was able to classify 51% of the GPC OTUs at the genus level, which is almost double the classification rate of GreenGenes2 (**Figure 2e**).

The MFD FL18S rRNA gene sequences revealed eukaryotic diversity across the different habitats (**Figure 2f**). The rRNA operon sequences exhibit a strong phylogenetic signal as they include both the ITS1 and ITS2 regions. However, the absence of a comprehensive rRNA operon reference database prompted us to focus our analysis on the FL18S rRNA genes that can be directly compared to the PR2 database^26^. The 13.4 million eukaryotic FL18S rRNA gene sequences clustered into 12,440 species (99% sequence identity, **Supplementary Note 1**). Mapping of the species-representative sequences against PR2 revealed that most species (77%) are novel (**Figure 2b**). Furthermore, 32% of the sequences had <93% similarity to a sequence in PR2, indicating high novelty at approximately the family level (**Supplementary Note 1**). These findings underscore that vast micro-eukaryotic diversity remains undocumented.

### Alpha and gamma diversity as habitat management discriminators

The level of microbial diversity in a habitat is often characterised by the alpha diversity, the richness in a single sample or average sample of a habitat, and by the gamma diversity, the total observed richness of all samples within a habitat^30^. In contrast to the above ground macro biodiversity, it has been shown that disturbed, managed soils have higher richness than undisturbed natural areas, both at continental and global scales^30,31^. Our detailed ontology and the number of samples in each habitat type enabled us to reevaluate these observations using the rRNA gene datasets.

The highest bacterial alpha diversity was observed in the disturbed or highly disturbed areas, with “Greenspaces’’ having a ∼3-fold larger observed richness than “Forests” (p<0.001) (**Figure 3a**), supporting high species richness in disturbed areas^30,31^. However, the prokaryotic gamma diversity was significantly lower in highly disturbed areas such as “Fields” [N=93, 95% CI: 18,266-18,487], compared to less disturbed areas, e.g. “Forests” [N=75, 95% CI: 27,819-28,315] (**Figure 3a**, **Supplementary Note 2**). This effect was most pronounced among prokaryotic communities **(Figure 3a**) but also visible in the eukaryotic data (**Supplementary Figure 3**) and agreed with the above ground macro biodiversity^32^. These results show that the more disturbed habitats harboured the largest number of different species in a single sample (or average sample), but with greater species homogeneity as consequence of many shared species across the habitat (lower within group dissimilarity, supported by the Bray-Curtis analysis **Figure 3a**) as recently noted by other studies^33,34^. This suggests that gamma diversity should be included as a metric for monitoring microbial diversity across habitats, e.g. when evaluating effects of land use perturbation. Furthermore, it highlights the need for more large-scale studies at finer spatial resolution to carry out gamma diversity analyses.

**Figure 3:**
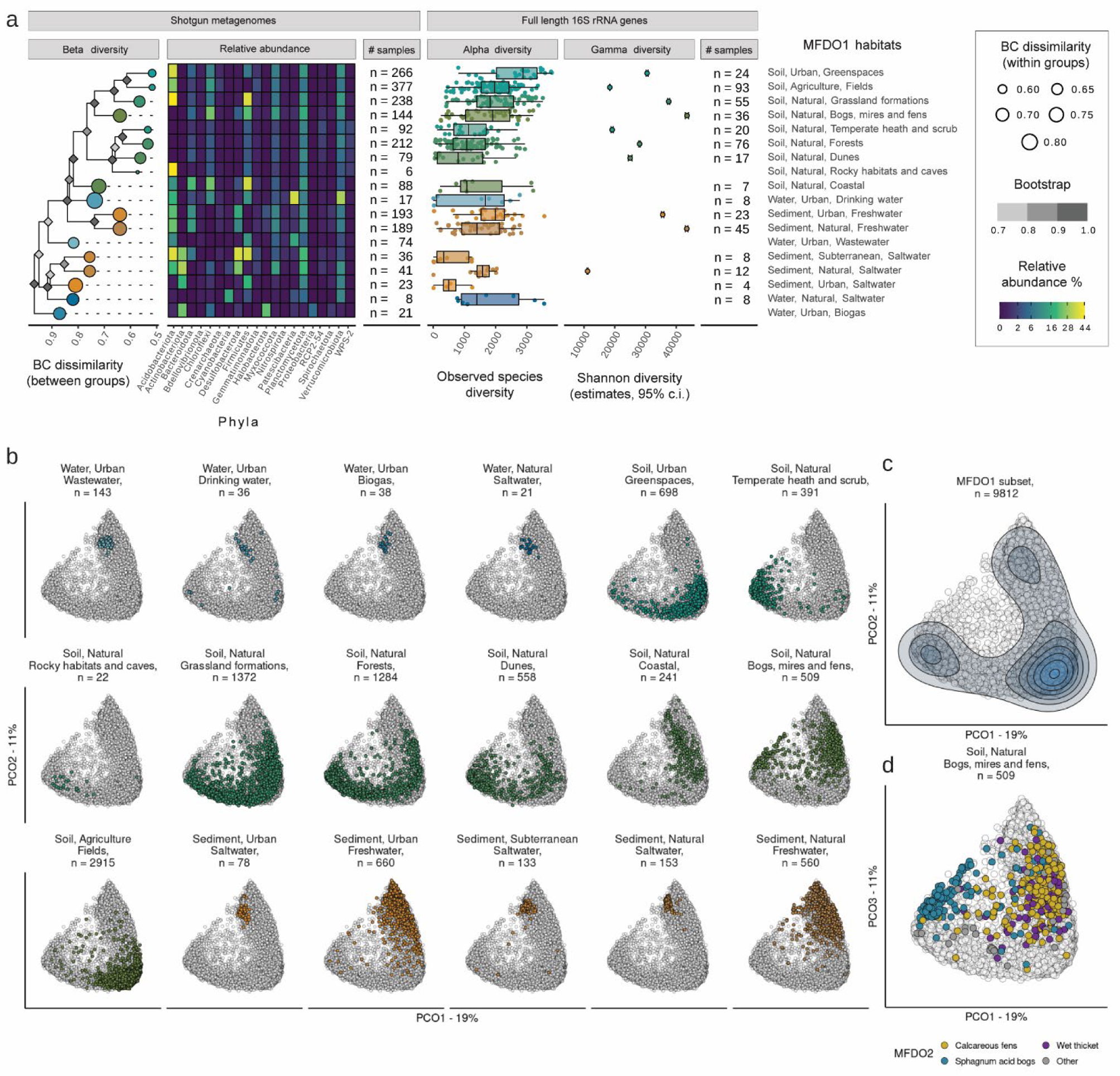
Microbial diversity of the Danish habitats. **a.** Ordination of the full metagenome dataset. The 10,686 metagenome samples are arranged using PCoA and the 18 selected MFDO1 habitats are highlighted in different panels and colours. The samples of each habitat often occupy a distinct portion (of variable size) of the plot. **b.** The contour plot depicts the density of points in the ordinations space. **c.** The sample distribution in the ordination space for the MFDO1 “Soil, Natural, Bogs, Mires and Fens” with classifications at the MFDO2 ontology level. **d.** Diversity overview of the habitats. The metagenomes represent the samples from the reference grid. The relative abundance of the 20 most abundant phyla are shown. Bray Curtis (BC) dissimilarity. Bootstrap values were calculated using 500 iterations. The category “Rocky habitats and caves” includes only 6 samples from the island of Bornholm, which is geologically different from the rest of Denmark (**Supplementary Figure 2**). The three hinges of the box plots correspond to 25^th^, 50^th^ and 75^th^ percentiles of the distributions whilst the whiskers extend to a maximum of 1.5 times the distance between the 25^th^ and 75^th^ percentile hinges. All individual samples are shown as points (with a jitter to improve visualisation). Average Shannon diversity is reported with corresponding error bars spanning the 5^th^-95^th^ percentiles of the distribution. Gamma diversity values are depicted only for habitats with at least 10 samples.

### Convergence of supervised and unsupervised habitat descriptors

The Microflora Danica dataset presents a unique opportunity to investigate how microbial communities corroborate habitat ontologies defined by macroflora and abiotic observations. Generally, we found a good separation between the different MFDO1 habitats in ordination space using the 16S rRNA gene genus-level microbial community composition of 9,812 samples (**Figure 3b,c**). An exception is MFDO1 “Bogs, mires and fens”, which showed large dispersion in ordination space. At the MFDO2 level, this habitat class consists of both “Calcareous fens” and “Sphagnum acid bogs”, which have large differences in pH that impact the microbial communities^35^ **(Figure 3d)**. To further substantiate the link between microbial communities and habitat ontologies we first estimated spatial autocorrelation using distance decay analysis (**Supplementary Figure 4**) and, based on this, identified representative samples of the MFDO1 habitats within the 10 km reference grid of Denmark (2,122 samples, **Supplementary Figure 2**, see **Methods**). Using the spatially thinned dataset, we calculated within-group and between-group Bray-Curtis dissimilarities across all combinations of samples. At the high-resolution country-scale of Microflora Danica, sample clustering largely captured the expected relationship between habitats at the MFDO1 level, confirming previous global observations^5,36^ (**Figure 3a**).

To take advantage of the high-resolution MFD ontology, we used the prokaryotic 16S rRNA gene fragment counts as predictor variables in a random forest habitat classification across all MFD ontology levels and taxonomic ranks (phylum to genus) (**Figure 4**, **Supplementary Note 3**). Some habitats were difficult to model, e.g. all agricultural fields, where the level of shared taxa was large, while habitats with highest model scores were more specialised microbiomes (e.g. “Saltwater” and “Wastewater”). In general, low model scores reflected habitats where samples would be misclassified to a few other selected habitats e.g. samples from “Grassland formations”, “Greenspaces” and “Fields” were often misclassified as each other (**Supplementary Note 3**). Considering which prokaryotic orders were the most important in discriminating among habitats, the strongest signal was provided by Solirubrobacteriales, whose species have been found in soils and roots, but accounted for the largest relative abundance in deserts^37^. Coherently with literature, relative abundance of Solirubrobacterales showed a peak in “Fixed dunes” and “Xeric sand calcareous grasslands”, whilst being present, to a minor extent, in other habitats and, perhaps, functioning as a general proxy for sand content in the samples. Our findings support the proposal of defining habitats based on continuous gradients instead of discrete categories^38–40^, and that microbiome data might be an important datasource that could be a scalable solution to future classification efforts.

**Figure 4:**
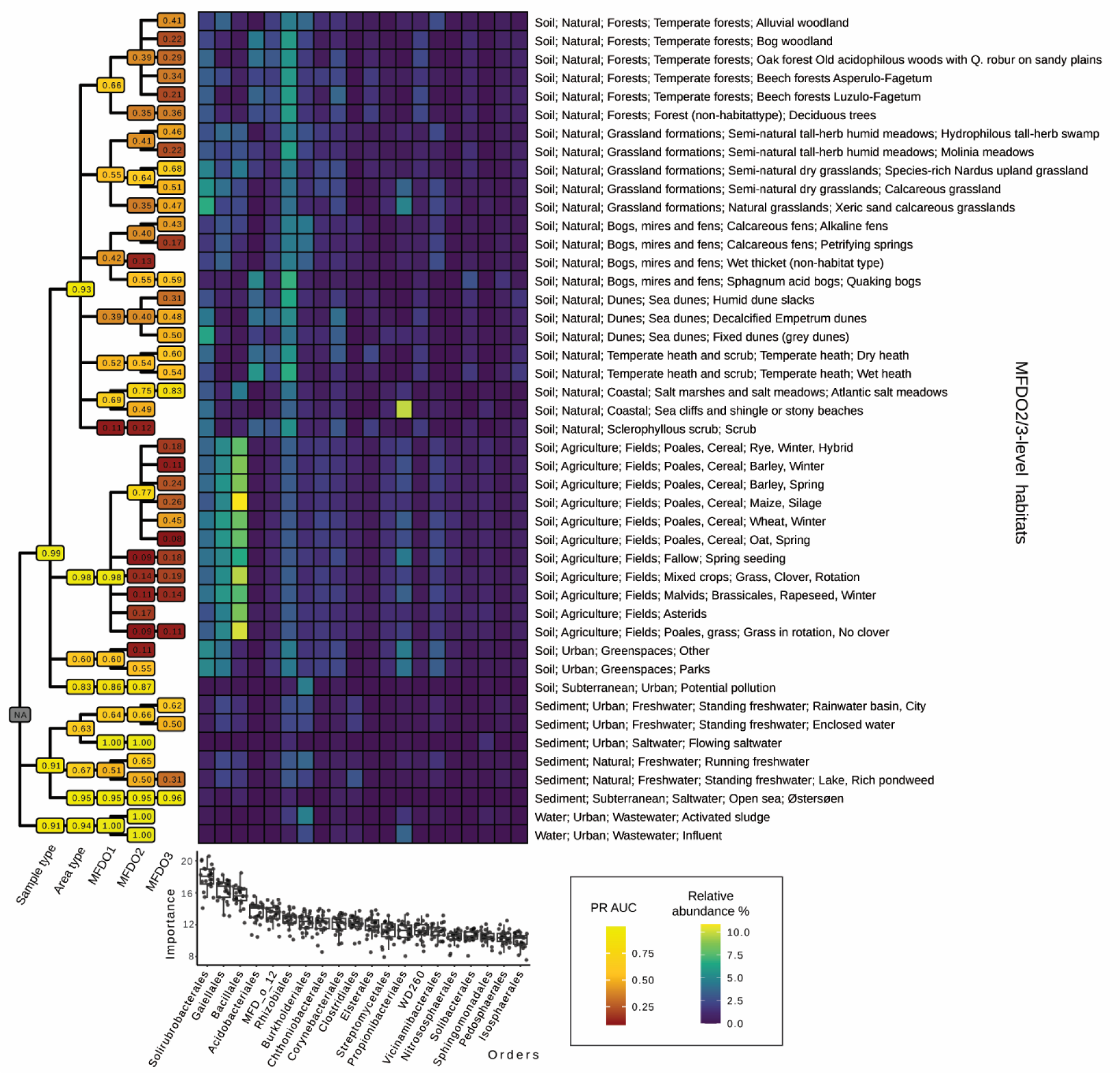
Per-class assessment at the order level. The Order-level models were used to compile the per-class Precision Recall Area Under the Curve (PR AUC) of every node of the ontology. The mean results (n=25: 5 folds x 5 iterations) are reported in the tree labels and coloured accordingly, with brighter nodes carrying higher values. Moreover, the top 20 Orders, according to variable importance (boxplot on the bottom, computed using the MFDO3 models), are reported with their median relative abundance for each of the terminal nodes of the ontology (at the MFDO3 or MFDO2 level). Of note, the models were reliable in classifying samples from agricultural soils (PR AUC=0.93) but not at classifying individual crop types.

### Core genera are abundant and prevalent across habitats in Denmark

We identified abundant core community genera in the habitats at all ontology levels (mean relative abundance >0.1% and >50% habitat-specific prevalence, **Supplementary Table 1**). With increasing specificity of the ontology, the median size of the core community increased from 28 to 55 genera and 13% to 42% relative abundance. Across all MFDO1 habitats 324 core genera were identified (1.5% of the total 20,961 identified genera) comprising 29% of the relative abundance of classified 16S rRNA gene fragments (**Supplementary Figure 5**). This high classification fraction aligns with patterns described across terrestrial ecosystems of Earth, where the dominant phylotypes (2% of the total) constituted an average 41% of the soil bacterial communities in terms of relative abundance^41^.

Across all MFD habitat categories, 565 core genera were identified. The 10 most prevalent core genera across the 57 MFDO3 habitats were *Mycobacterium* (prevalence: 50, median relative abundance: 1.8%), *Bacillus* (45, 2.3%), *Candidatus* Solibacter (44, 0.8%), *Haliangium* (44, 0.7%), *Bryobacter* (44, 0.5%), *Conexibacter* (43, 1.0%), *Paenibacillus* (43, 0.4%), *Pajaroellobacter* (42, 0.3%), *Bradyrhizobium* (41, 0.4%) and one genus with MFD placeholder name, MFD_g_8 (40, 0.5%) from the family of *Xanthobacteraceae* (**Supplementary Table 2**). Outside the top 10, *Acidothermus* (30, 7.3%) stands out, with a relative abundance of up to 21%, as observed in the temperate dry heaths of Denmark (N=52).

Habitat-specific core genera, or indicator taxa, were mostly identified in habitats with strong selecting environmental gradients (e.g. halotolerance) or well-defined habitats such as biogas systems (**Supplementary Table 1**), suggesting a high level of microbial prevalence if no unique environmental selective pressure is present and underlines that the prokaryotic community aligns across continuous gradients. In soil, for example, the median of core genera unique to a specific MFDO1 habitat was 2 (**Supplementary Table 1**), showing that many of the genera were shared among two or more habitats. This was further underlined by the lack of indicator taxa for the differentiation of crops on agricultural fields (median 0), and also reflected in a small difference in the core-genera size between the area type agriculture (67) and MFDO3 specific crop type (72) (**Supplementary Table 1**).

### MAGs from Danish habitats double the known species fraction in metagenomes

The MFG 16S rRNA gene database is a comprehensive resource for genus-level taxonomic investigations of microbial communities, but our analysis revealed a large shared core microbial community at this level of taxonomic resolution. A larger difference between habitats might be identified on genome-level, but the quality of such analysis relies heavily on the representativeness of the genome databases. Therefore, we used SingleM^42^ to estimate the fraction of known species in our 10,686 MFD metagenomes when compared to a collection of 167,824 species-level genomes and MAGs (95% ANI dereplicated) compiled from all major public databases (GTDB-R214^10^, SPIRE^8^, GEM^7^, SMAG^11^, Ocean MAGs^12^, and UHGG^9^).

Compared to 30,699 soil, sediment and water metagenomes in NCBI SRA (downloaded Dec 2021^42^), the 10,686 MFD metagenomes contained the least amount of known species. At the MFD sample type level, the mean percentage of known species in the MFD metagenomes was lowest in soils (6.4±3.7%), followed closely by sediment (7.0±6.0%). Biogas and wastewater treatment plants had the highest proportion of known species at the MFDO1 level (mean 75.6% and 39.5%, respectively), confirming that human-associated environments are better sampled and characterised than natural environments^2,43^ (**Figure 5a, Supplementary Table 3**). Interestingly, the metagenomes also contained high proportions of non-prokaryotic DNA (14-43% of reads, **Supplementary Figure 6**) that could be a key resource for future analyses^44^.

**Figure 5:**
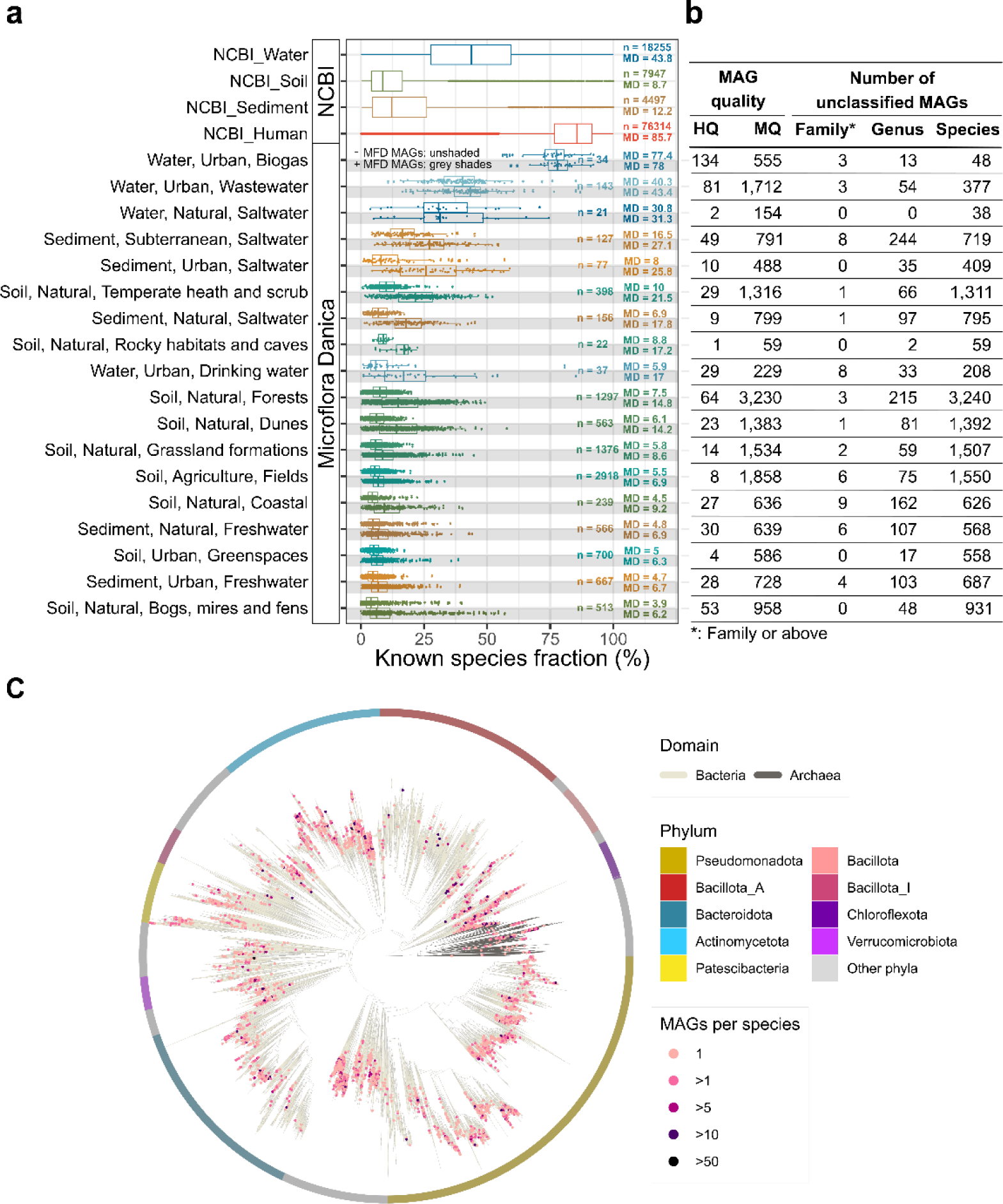
Novelty of Danish microbes. **a.** The known species fraction of the microbial community in each metagenome estimated using SingleM against 167,284 MAGs from public catalogues (unshaded, see **Methods**) and with the MFD MAGs included (grey-shaded). The first 4 bars represent 30,699 public metagenomes downloaded from NCBI SRA. **b.** MAG recovery statistics and taxonomic novelty against GTDB-R214. **c.** Genome tree of the 5,518 species representatives MAGs recovered from the MFD metagenomes.

The *de novo* assembly of the 10,686 metagenomes recovered 19,253 bacterial and archaeal MAGs that belonged to 15,791 new species and 1,507 new genera (**Figure 5b**), with broad phylogenetic coverage (**Figure 5c**). Adding these MFD genomes to the collection of public genomes almost doubled the amount of known species that could be identified in the MFD metagenomes using SingleM (**Figure 5a**, **Supplementary Table 4**). However, despite the large progress, the recovered MAGs only cover 8.2%-31.8% of the prokaryotic communities (**Figure 5a**, **Supplementary Figure 7**), underlining that more large-scale sampling and sequencing efforts are needed to obtain representative genomes across the tree of life, especially in soil and sediment samples. This is essential for investigating organisms with disproportionately large impacts on our global processes, such as those involved in the nitrogen cycle.

### Distribution patterns of canonical and novel nitrifiers across Danish habitats

Soil fertilisation with reactive nitrogen is essential to feed about 50% of the human population^45^, but is at the same time the major source of nitrogen pollution globally^46–48^. Nitrification, the aerobic conversion of ammonium to nitrite and nitrate, is an essential step in the biogeochemical nitrogen cycle and contributes to massive nitrogen fertiliser loss in agriculture and the production of the potent greenhouse gas and ozone-depleting substance nitrous oxide^49,50^. The microbial functional groups responsible for nitrification^51,52^ are ammonia oxidising bacteria (AOB) and archaea (AOA), nitrite oxidising bacteria (NOB) and complete ammonia oxidising bacteria (CMX).

In order to investigate the diversity, distribution and novelty of nitrifiers across Denmark, we improved gene-based search models by incorporating the nitrification marker genes *amoA* (encoding a subunit of the ammonia monooxygenase of AOB, AOA and CMX) and *nxrA* (encoding the active-site subunit of nitrite oxidoreductase of NOB and CMX), from the recovered MFD MAGs (**Figure 6b**, **Supplementary Note 5**). Distinct differences in the nitrifier communities based on gene abundances were observed between MFDO1 habitats (**Figure 6a**), though we identified similar profiles between the following pairs of habitats: natural and urban sediments, fields and greenspaces, and forests and grasslands. The nitrifier community showed similar ordination patterns to the entire microbial community on the MFDO1 habitat level (**Figure 3a**). In addition, the nitrifier community showed strong separation of MFDO2 habitats types, highlighted by the splits between “Sphagnum acid bogs” and “Calcareous fens”, “Alluvial woodland” and “Non-native- and Deciduous trees”, and “Semi-natural dry grasslands” and “Semi-natural tall-herb humid meadows” (**Figure 6a**).

**Figure 6:**
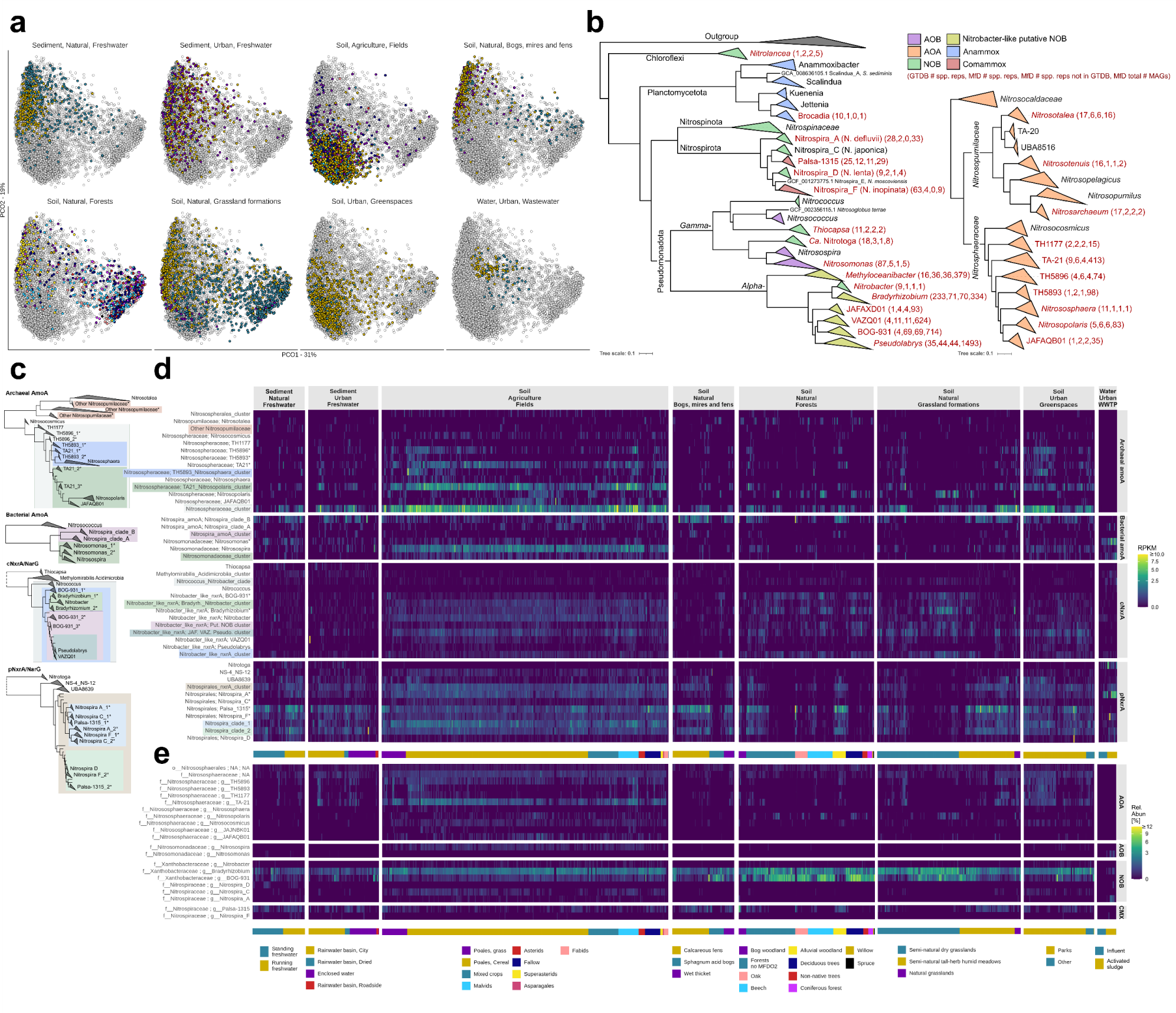
Nitrifier distribution in Danish Habitats. **a.** Ordination of functional nitrifier genes. Samples from 8 selected MFDO1 habitats are arranged using PCoA based on nitrifier gene distribution. Samples classified with a MFDO2 containing ≤ 2 samples were not included. The 8 selected MFDO1 habitats are highlighted in different facets and coloured by MFDO2, as depicted in **e**. The visualisation depicts the first two components, accounting for 31% and 19% of explained variance, respectively. **b.** Phylogenomic tree of nitrifiers. The red text indicates groups for which we recovered MAGs, with the numbers in the brackets indicating number of species in GTDB, number of species recovered in MFD, number of species recovered in MFD not present in GTDB, and total number of MAGs recovered in MFD**. c.** Phylogenetic trees of enzymes essential for nitrification. Protein phylogeny of archaeal AmoA, bacterial AmoA, cytoplasmic NxrA/NarG, and periplasmic NxrA/NarG. Clades denoted by * holds sequences from a genus separated into several clades that have been combined in Figure 6d. Groups encompassing several clades are shaded in corresponding colours to Figure 6d. **d.** Distribution of nitrification genes across Danish habitats. Number of reads (RPKM, Reads Per Kilobase Million) assigned to each gene-phylogenetic group displayed in 6c. Samples are clustered with hierarchical clusters within each MFDO2 habitat. A bottom panel indicates the MFDO2 habitat. **e.** Distribution of canonical and potential nitrifiers across Danish habitats based on SingleM. Heatmap of short-read metagenomes based on SingleM with metapackage supplemented with MAGs from short-read metagenomes.

The highest relative gene abundances of canonical ammonia oxidizers (AOA and AOB) were observed in “Fields” and “Greenspaces” (**Figure 6d**). These habitats displayed similar nitrifier communities, and were dominated by AOA, particularly those belonging to the so far uncharacterised genera TA-21 and TH5896 (*Nitrosophaeraceae*), from which we recovered many MAGs (**Figure 6b**). Their high abundances were supported by profiles using SingleM analysis and the detection of marker genes from genomes (**Figure 6d,e**). Moreover, the *de novo* genus MFD_g_4907 (*Nitrosophaeraceae*) appeared as a shared core genus between “Fields” and “Greenspaces” (**Supplementary Note 4**). In agreement with findings of *Nitrosospira* and AOA being prevalent in agricultural soils^53–55^, we found that *Nitrosospira amoA* was abundant compared to other AOBs, but in lower relative abundance than *Nitrososphaeraceae* AOAs.

Recent studies, however, have challenged the dominant role of AOA and canonical AOB in soil nitrification processes, suggesting that CMX *Nitrospira* may be more abundant^56–59^. *Nitrospira* was present across all MFDO1 levels investigated based on 16S rRNA gene fragments. Canonical and CMX *Nitrospira* are difficult to differentiate based on *nxrA* and 16S rRNA genes^60^, but by using *amoA* gene-phylogeny CMX *Nitrospira* can be identified and divided into clades A and B^56^. These two clades are found within the genera Nitrospira_F and Palsa-1315 (GTDB v214), respectively. Palsa-1315, first named from MAGs found in a permafrost peatland^61^, is likely a new comammox containing genus. This is supported by a linear correlation [R^2^ = 0.34-0.77] between *Nitrospira* clade B *amoA* and Palsa-1315 *nxrA* in various MFDO1 habitats (Supplementary Figure 8) resolved by our improved search database (**Figure 6d**), and was corroborated by SingleM results (**Figure 6e**).

Interestingly, our improved search models showed that CMX clade B was more abundant than CMX clade A in most habitats, especially natural soils and sediments, and in particular MFDO2 habitats “Calcareous fens”, “Alluvial woodland”, and “Semi-natural humid meadows’’ (**Figure 6d,e**), likely driving the observed separation (**Figure 6a**). This challenges the perception that CMX clade B is not abundant in forest soils^62^, wetland sediments^63^, and acidic or fertilised agricultural soils^58,59,64^.

*Nitrospira* clade A *amoA* was inconsistently identified in sediments and agricultural soils, and nearly absent from the other habitats investigated (**Figure 6d**). Our analysis highlights CMX clade B as the most abundant ammonia oxidiser in natural habitats, while canonical AOB and AOAs were more abundant in managed habitats. In fields, a substantial part of *Nitrospira-*like *nxrA* sequences were ascribed to canonical *Nitrospira* NOB, but unresolved at genus level. These were likely *Nitrospira C* based on single copy protein analysis (**Figure 6d,e**).

*Nitrobacter* is considered to be an abundant NOB in fertilised soil, based on *nxrA* identification^65,66^. However, our detailed search for NXR encoding MAGs indicates that *Nitrobacter* previously might have been over-classified, as we found a large diversity of NxrA sequences (>600 aa) falling between *Nitrobacter* and *Nitrococcus* unassociated with known NOBs and representing either uncharacterized NOBs or denitrifiers (**Supplementary Note 5**). While other studies have reported cytoplasmic *nxrA* sequences clustering near, but outside of, cultivated *Nitrobacter* representatives in agricultural soils^66^, we have linked these *Nitrobacter*-like NxrA sequences to members of the *Xanthobacteraceae* family, primarily *Bradyrhizobium* spp., *Pseudolabrys* spp. and the uncharacterised genus BOG-931 (**Figure 6b**, **Supplementary Note 6**). In particular, a monophyletic clade of *Nitrobacter*-like NxrA from BOG-931 grouped close to *Nitrobacter* and *Nitrococcus* NxrA, while genomes containing these sequences also cluster together in a phylogenomic tree (**Supplementary Note 6**). Inspecting gene synteny of the *nxr/nar* operons and other relevant genes strongly indicates that members of BOG-931 could be potential new NOBs, but confirmation requires culturing (**Supplementary Note 6**).

The putative *Nitrobacter*-like *nxrA* groups were found across fields, forests, grasslands and greenspaces (**Figure 6d**). In fields, *Nitrobacter* and *Nitrobacter*-like *nxrA* genes were present independent of crop type, and were less abundant than canonical *Nitrospira* NOB (**Figure 6d**). In line with this, *Nitrobacter* and *Nitrobacter*-like *nxrA* groups have been linked to high N availability in agricultural soils^66^. BOG-931 was most abundant in forest soils, and *Nitrobacter* and *Nitrobacter*-like NOBs have previously been associated with nitrogen amendment in forest soils^67^ and alpine meadows^68^. Indeed, BOG-931 appeared to be detected mainly in soil habitats lacking detected CMX clade B, especially pronounced in forests, grasslands and acid bogs of the “Bogs, mires and fens” (**Figure 6d**). This suggests niche differentiation between CMX and canonical NOBs, and underlines a general need for further investigation of novel and often abundant nitrifiers, including *Nitrobacter nxrA*-like groups, CMX clade B, and AOAs TA-21 and TH5896. As the presence and abundance of nitrifiers may be applied to evaluate how human activities affect the nitrogen cycle^53,64^, our results stress the importance of developing and applying reliable methods for diversity and distribution analysis, to include all important groups, including the novel nitrifiers, and to understand their response to environmental factors.

## Conclusions

Here, we provide a census of Denmark’s free-living environmental microbial communities, and the baseline of a nation’s microbial diversity by investigating 10,686 samples across the major habitat types in Denmark. Using 449 representative samples we generated more than 30 million rRNA operon sequences from bacteria and eukaryotes, expanding the current databases by an order of magnitude, to create the Microflora Danica rRNA reference database. Using the rRNA data we estimate approximately 75,000 common bacterial species, which serve as a lower bound for targeted efforts to understand the species that likely are important for the main ecosystem functions. Previous large-scale microbial studies have lacked resolution in habitat classification, greatly limiting the scope of use. In Microflora Danica all sampled natural habitats are classified by domain experts according to the highly specific EUNIS and Natura2000 classification schemes based on macroflora and geographic location. To encompass all habitats sampled in Microflora Danica and previous high-level microbial habitat classification schemes (EMPO) we generated the 5-level Microflora Danica Ontology. Leveraging the MFD samples, rRNA database and ontology enabled us to conduct analysis of 10,686 shotgun metagenomes at unprecedented geographic, taxonomic and habitat resolution. While many habitats have distinct microbial profiles, we identified several habitat types with similarities that could not be identified using above ground classification based on macroflora. These similarities may be explained by histories of active land management and disturbances, which can increase species diversity, but also increase homogeneity as the communities in the managed environments become increasingly similar. This pattern is also recreated at the functional level, with nitrifier communities highly reflective of the habitat types and level of human disturbance. We determine that biodiversity assessments and monitoring need to accommodate gamma diversity information to prevent homogenisation of the Danish microbiome. Future assessments could be conducted using a data-driven approach, as we show that models can use short-read data to match a microbiome to flora-inferred habitats. Our comprehensive data enabled us to investigate the diversity and distribution of canonical and novel nitrifiers across habitats, including natural, agricultural and urban areas. Surprisingly, the results strongly challenge existing perceptions concerning prevalent genera and their habitat preference and also reveal novel nitrifiers as more abundant than canonical nitrifiers. This suggests niche differentiation between CMX and canonical NOBs, and underlines a general need for further investigation of novel and often abundant nitrifiers, including *Nitrobacter nxrA*-like groups, CMX clade B and AOAs TA-21 and TH5896. As new nitrifiers are identified and characterised, informed control and management of agricultural practices based on nitrifier populations might be within our reach by linking specific populations to land use, fertiliser regimes, and greenhouse gas emissions. The Microflora Danica datasets provide a valuable resource for future genome mining efforts, for comparisons against international samples and as a reference point to monitor future changes within Denmark’s microbial community.

## Data availability

The sequencing data, and metagenome bins (>90 % completeness <5 % contamination) are available in NCBI under bioproject PRJNA1071982. Metadata and datafiles are available on github (https://github.com/cmc-aau/mfd_wiki). Additional datafiles (metagenome assemblies, metagenome bins) are present in Zenodo DOI: 10.5281/zenodo.10695834. The MFD and MFG databases are available on figshare: 10.6084/m9.figshare.26030524.

## Acknowledgements

Funding was provided by the Poul Due Jensen Foundation, PDJF (grant MicroFlora Danica to MA and PHN), Villum Foundation (grant 15510 and 50093 to MA, grant 13351 to PHN), the European Union (ERC grant 101078234 to MA). We want to thank the Microflora Danica Consortium for their invaluable contribution by collecting samples and related metadata across Denmark.

## The Microflora Danica Consortium

Henning C. Thomsen^1^, Bent T. Christensen^1^, Lis W. de Jonge^1^, Anne-Cathrine S. Danielsen^1,2^, Cecilie Hermansen^1^, Mogens H. Greve^1^, Rasmus Ejrnæs^3^, Thomas Davidson^3^, Signe Normand^2,4^, Urs Treier^2,4^, Bjarke Madsen^2,4^, Andreas Schramm^5^, Ian Marshall^5^, Kasper U. Kjeldsen^6^, Kai Finster^7^, Philip F. Thomsen^7^, Eva E. Sigsgaard^7^, Martin J. Klepke^7^, Marie Vestergård^7^, Erik Aude^8^, Lene Thomsen^8^, Camilla Lemming^9^, Rita Hørfarter^9^, Marlene M. Jensen^10^, Tobias G. Frøslev^11^, Lone Gram^11,12^, Peter B. Svendsen^11,12^, Morten Dencker Schostag^11,12^, Sanne Kjellerup^13^, Torben L. Skovhus^14^, Ditte A. Søborg^14^, Kasper R. Jensen^15^, Jørgen F. Pedersen^16^, Andrew Giguere^17^, Inge S. Pedersen^18,19^, Mads Sønderkær^18,19^, Jes Vollertsen^20^, Fan Liu^20^, Peter Roslev^21^, Niels Iversen^21^, Kåre L. Nielsen^21^, Nadieh de Jonge^21^, Dan Bruhn^21^, Jeppe L. Nielsen^21^, Torsten N. Kristensen^21^, Chenjing Jiang^21^, Marta A. Nierychlo^21^, Giulia Dottorini^21^.

^1^Department of Agroecology, Aarhus University, Tjele, Denmark. ^2^Center for Sustainable Landscapes under Global Change, Aarhus University, Aarhus, Denmark. ^3^Department of Ecoscience, Aarhus University, Aarhus, Denmark. ^4^Section for Ecoinformatics & Biodiversity, Department of Biology, Aarhus University, Aarhus, Denmark. ^5^Center for Electromicrobiology, Department of Biology, Aarhus University, Aarhus, Denmark. ^6^Section for Microbiology, Department of Biology, Aarhus University, Aarhus, Denmark. ^7^Department of Biology, Aarhus University, Aarhus, Denmark. ^8^Habitatvision, Lystrup, Denmark. ^9^SEGES Innovation P/S, Aarhus, Denmark. ^10^Department of Environmental and Resource Engineering, Technical University of Denmark, Kgs Lyngby, Denmark. ^11^Global Biodiversity Information Facility, Denmark. ^11^Department of Biotechnology and Biomedicine, Technical University of Denmark, Kgs. Lyngby, Denmark. ^12^Center for Microbial Secondary Metabolites, Technical University of Denmark, Kgs. Lyngby, Denmark. ^13^WSP Danmark A/S, Tåstrup, Denmark. ^14^Research centre for built environment, climate and water technology, VIA University College, Horsens, Denmark. ^15^Department of Biology, University of Southern Denmark, Odense, Denmark. ^16^Skalhuse, Nibe, Denmark. ^17^Centre for Microbiology and Environmental Systems Science, University of Vienna, Vienna, Austria. ^18^Department of Molecular Diagnostics, Aalborg University Hospital, Aalborg, Denmark. ^19^Department of Clinical Medicine, Aalborg University, Aalborg, Denmark. ^20^Department of the Built Environment, Aalborg University, Aalborg, Denmark. ^21^Department of Chemistry and Bioscience, Aalborg University, Aalborg, Denmark.

## Materials and Methods

### Sampling

The samples include those collected as part of the Microflora Danica sampling campaign or contributed by collaborators. MFD samples were registered and associated with the appropriate meta using codeREADr [https://www.codereadr.com] using a linear barcode attached to sterile 100 mL sample containers. After collection the samples were stored between 4°C and 10°C before being deposited at -20°C for later processing. Up to five topsoil subsamples (0-20 cm), taken within a ∼80 m^2^ (5 m radius) sampling area, were mixed in a sterile plastic bag and transferred to the sample containers (P04_2, P04_4, P04_6, P04_7, P08_1, P08_2, P08_3, P08_5, P08_6, P08_7, P08_8, P17_1). For subterranean soils different samples were collected at different depths (P06_1, P06_2, P06_3). Soil samples provided by Aarhus University and Copenhagen University were collected as described in Brunbjerg et al. 2019^39^. Briefly, 81 subsamples, spanning a 9 x 9 grid covering a 40 x 40 m plot, were mixed into a representative sample (P01_1, P01_2, P02_1, P02_). Samples from Land Use and Coverage Area frame Survey (LUCAS, P04_8) were collected as described in Labouyrie et al. 2023^31^. Dried soils from croplands were provided as dried soils by SEGES. Individual samples were frozen, crushed to particles below 1 cm in size, and air dried at 37°C (P04_3, P04_5). These samples were rehydrated using phosphate-buffered saline before downstream processing. Other soil samples from natural and agricultural land were collected at a single point (P03_1^69^, P04_1, P19_1, P21_1).

Up to five sediment subsamples (0-10 cm) across the sampling area were mixed in a sterile plastic bag after careful manual removal of any collected water and transferred to the sample container (P05_1, P05_2, P09_1, P11_1, P21_1). For sediment cores collected from streams, the top layer (0-5 cm) was sampled in MFD (P10_1, P10_2, P10_3). These samples were collected as three subsamples across a 20 m transect of the stream, two at 25% distance from each brim and one in the middle of the stream. Sediments from fjords were collected together with water, and filtered onto a dead-end filter (P11_3). Harbour samples were collected as scrape offs from different surfaces (fenders, piers, etc.) with three samples from each location (P18_1, P18_2). Sediment samples provided by University of Southern Denmark were collected from the deepest part of the lake as described in Hassler et al. 2024^70^ (P09_3, P09_4). Other sediment samples from coastal areas were collected at a single point (P11_2^71^, P12_3). Surface and subterranean sediments from the Baltic sea were provided by the Danish Environmental Agency (P12_5). These subterranean sediments were collected using a Vibrocore sampler. Other surface and subterranean sediments were collected by gravity corer, haps corer, and Rumohr corer (P12_1, P12_2).

Samples of drinking water from the water treatment plants were filtered onto dead-end filters (P16_1, P16_2, P16_3, P16_4). Samples of sand filter material were also included.

Samples from specific sampling projects were provided as DNA extracts and included samples from polluted soil and groundwater (P06_3), coastal and fjord sediments (P11_2)^71^, and marine sediments (P12_1, P12_2)^72–75^. Wastewater was sampled as both influent (P13_1), using flow proportional sampling, and activated sludge from the aeration tank (P13_2) as described in Dottorini et al. 2021^76^. Sludge was also collected from anaerobic digesters (P13_3) as described in Dueholm et al. 2024^19^.

The number of subsamples and other related metadata can be found on github.

### MFD ontology

The MFD Ontology (MFDO) was developed as a link between the classical plant-derived habitat ontologies and the Earth Microbiome Project Ontology (EMPO). The broadest MFDO classification level (sample type), corresponds to the most specific EMPO level (EMPO level 4^6^), while the detailed levels for natural samples correspond broadly to the Natura 2000 habitat ontology ^15^. Finally, missing categories, such as urban, were adapted from the EUNIS ontology^16^ to provide a detailed description of non-natural habitats. The MFDO was designed -for the moment -to fit the Danish environment and it was refined with a panel of national experts. The full MFDO and its association to other habitat ontologies (i.e. EMPO, Natura 2000 and EUNIS) can be found at https://github.com/cmc-aau/mfd_metadata.

### Metadata curation

The metadata collected with codeREADr was screened for completeness in the following fields (hereafter addressed as “minimal metadata”): fieldsample_barcode (the unique sample identifier), project_id (unique identifier of the subproject), longitude and latitude (ISO 6709), sitename (common name of the sampling site), coords_reliable (indicating if the coordinates are reliable, not reliable or masked), sampling_date (sampling date ISO 8601) and five levels of the MFD habitat ontology (mfd_sampletype, mfd_areatype, mfd_hab1, mfd_hab2, mfd_hab3). If a sample presented an incorrect entry, the error was corrected using R^77^ v4.2.3, and if a correction was not possible, the responsible person for the subproject was contacted. The process was iterated until improvements were not anymore possible. Briefly, common corrections included case changing, date formatting (using lubridate^78^ v1.9.2) and coordinate projection (*project* function form terra v1.7.55). The reference grid mapping and masking of the coordinates were performed using the terra^79^ v1.7.55 package. The European Environment Agency 1-km^80^ and 10-km^81^ reference grids of Denmark were projected from EPSG:3035 to EPSG:4326 (function *project*), whilst the coordinates from MFD samples were encoded into a spatial vector (function *vect*) and mapped on the grids (function *intersect*) to identified their cells of origin. The cells associated with each sample (when the coordinates were present), were reported in the fields “cell.1km” and “cell.10km” of the metadata. The centroids of the cells were computed (function *centroids*) and, in case of samples from subprojects P04_3 and P04_5, the centroids were provided as latitude and longitude, whilst the coords_reliable field for those samples was set to “Masked”. The coordinates from subproject P06_3 were provided already masked as generic locations in the commune of sampling. Concordance between manual annotation of the habitat and government-registered LU (land use) designation was inferred comparing the MFDO for each sample with the Basemap04^82^ aggregated LU map. A broad correspondence of terms between MFDO and LU terms was established and, in order to account for GPS and labelling inaccuracies, any match in a range of 20 m was considered in concordance. Samples that were in disagreement were screened manually on Google Maps and if the disagreement was confirmed the coords_reliable field was set to “No”.

### Subsampling and DNA extraction

The sample containers with soil and sediment were thawed at 4°C and followed the procedure described in Jensen et al. 2024^83^ with the modification that the samples were divided into a total of three Matrix™ 1.2 mL 2D barcoded tubes [Thermo Scientific™]. Of the three 2D Matrix™ tubes only one was pre-filled with the lysing matrix E from MP biomedical, to which only 100 µL of sample material was added. The tubes destined for downstream DNA extraction were added to a 96-well SBS rack containing 4 empty positions, 4 reaction blanks and 1 extraction positive control. The layout can be found on github. Linkage of the linear barcodes of the original sample container, the 2D Matrix™ tubes and location in the final SBS racks was ensured with the use of a Mirage Rack Reader (Ziath) and the software DataPaq™ (Ziath) forwarding the entries to an SQL server (MongoDB). Pseudo-links were generated for samples received as DNA extracts.

The DNA extraction followed a slightly modified protocol of the DNeasy® 96 PowerSoil® Pro QIAcube® HT Kit [QIAGEN]. 500 µL CD1 was added to each 2D barcoded Matrix™ tube, whereafter the samples underwent three bead-beating cycles performed in two-minute intervals using the FastPrep96™ [MP™]. Between cycles the samples were kept on ice for two minutes. After lysis, samples were centrifuged at 3.486 x g for 10 minutes using an Eppendorf 5810 benchtop centrifuge. 300 µL supernatant was transferred by hand to a new S-block containing 300 µL CD2 and 100 µL nuclease-free water (NFW) to meet the requirement of 700 µL for the remaining part of the protocol. Samples were mixed by hand and centrifuged at 3.486 x g for 10 minutes, whereafter the sample transfer step was done using the QIAcube® HT [QIAGEN]. All subsequent steps followed the manufacturer’s protocol. Filters from dead-end filtration of water were extracted using the DNeasy® PowerWater® Kit [QIAGEN] according to the manufacturer’s protocol. The samples we received as DNA were previously extracted by our collaborators with either FastDNA™ Spin Kit for Soil [MP™] (polluted soil, influent wastewater, activated sludge and anaerobic digester sludge), DNeasy® PowerMax® Soil Kit [QIAGEN] (coastal sediments), or as a Phenol-Chloroform-Isoamylalcohol extraction (marine sediments). DNA was quantified using the Qubit 1X HS assay [Invitrogen™]. Extraction metadata including a denotation of methodology can be found on github.

### Short-read metagenomic library preparation, sequencing and processing

Metagenomic libraries were prepared with a 1:10 reagent volume reduction of the standard Illumina DNA prep protocol [Illumina] as described in Jensen et al. 2024^83^. Using the I.DOT One [DISPENDIX] 3 µL of up to 20 ng template DNA was prepared before addition of 2 µL BLT/TB1 and subsequent incubation in a thermocycler at 55°C for 10 minutes. Tagmentation was stopped by addition 1 µL TSB using the I.DOT One, and incubation in the thermocycler at 37°C for 15 minutes. The tagmented DNA was washed twice with 10 µL TWB. The I.DOT One was used to add the PCR master mix, prepared by mixing 2 µL EPM and 2 µL NFW per reaction. The epMotion® 96 [Eppendorf] was used to add 1 µL IDT® Illumina UD index to each reaction well. The input of genomic DNA was used to determine the applied cycles of the BLT-PCR program: 7 (4.9-20 ng), 8 (2.5-4.9 ng), 10 (0.9-2.5 ng), or 14 (<0.9 ng). Size-selection was performed on the libraries by addition of 17 µL NFW before 18 µL of the reaction volume was transferred to a new PCR-plate together with a mixture of 16:18 µL SPB:NFW. After incubation, 50 µL of the supernatant was transferred to a new PCR-plate with 6 µL undiluted SPB. After incubation, the beads were washed twice with 45 µL 80 % ethanol and eluted in 20 µL NFW.

The individual libraries were quantified using a 1:10 diluted upper standard. The pooled libraries were concentrated using 2x volume of SPRI ProNex® Chemistry [Promega] beads. Final sequencing libraries were produced by an equimolar combination of the pooled libraries. Quality control was performed using the Qubit 1X HS assay [Invitrogen™] and DS1000 or DS1000 HS ScreenTape [Agilent Technologies]. Library metadata can be found on github.

Metagenomic libraries were sequenced on the Illumina NovaSeq 6000 platform to a median depth of 5 Gbp. The Illumina data was demultiplexed using bcl2fastq2 v2.20.0. The raw reads were trimmed for barcodes, quality filtered, and deduplicated with fastp^84^ with the options” --detect_adapter_for_pe --correction --cut_right --cut_right_window_size 4 --cut_right_mean_quality 20 --average_qual 30 --length_required 100 --dedup --dup_calc_accuracy 6”. Commands were parallelized using GNU-parallel^85^ and outputs compressed using pigz^86^ v2.4. Sequencing metadata can be found on github.

### Full-length bacterial 16S rRNA gene amplicon library prep, sequencing and processing

A representative set of 415 samples were selected for 16S rRNA amplicon sequencing. Here, PCR was used to amplify the region V1-V8 of the 16S gene utilising UMI-tagged target primers enabling downstream chimera filtering and error-correction similarly to the method described by Karst et al. 2021^17^. All samples were tagged by the UMI-tailed target primers lu_16S_8F and lu_16S_1391R in a PCR reaction. The reaction contained 10-20 ng DNA input, 1X SuperFi buffer, 0.2 mM dNTPs, 500 nM of each primer, and 2 U of Platinum SuperFi DNA Polymerase [Thermo Fisher Scientific] in a total volume of 50 µL. The PCR-program consisted of initial denaturation at 95 °C for 2 minutes followed by 2 cycles of denaturation (95 °C for 30 seconds), annealing (55 °C for 1 minute) and extension (72 °C for 5 minutes). The PCR products were then purified with SPRI beads [CleanNGS] in a ratio of 0.7x bead/sample. Following 5 minutes of incubation the beads were washed twice in 80 % ethanol and eluted in 18 µL NFW for 5 minutes. The tagged molecules were then amplified in a second 25 cycle PCR reaction using barcoded primers targeting the UMI-adapter sequence. The PCR reaction contained 15 µL of the purified eluate, 1X SuperFi buffer, 0.2 mM dNTPs, 500 nM of forward and reverse primer, and 2 U of Platinum SuperFi DNA Polymerase [Thermo Fisher Scientific] in a total volume of 50 µL. The PCR-program consisted of initial denaturation at 95 °C for 2 minutes followed by 25 cycles of denaturation (95 °C for 15 seconds), annealing (60 °C for 30 seconds) and extension (72 °C for 3 minutes), followed by a final extension at 72° C for 5 minutes. The PCR products were purified with SPRI beads [CleanNGS] as described above and eluted in 20 µL NFW. Poorly performing samples underwent a third PCR reaction with 5-10 cycles using up to 50 ng amplicon DNA as input and otherwise identical to the previous PCR reaction.

The barcoded amplicons were multiplexed in pools of 5-6 samples containing a total of 300 ng. The pools were used as input for library preparation for DNA sequencing using the ‘Amplicons by Ligation (SQK-LSK110)’ protocol (version: ACDE_9110_v110_revG_10Nov2020) and loaded onto a MinION R.9.4.1 flow cell (FLO-MIN106D). The flow cells were sequenced for up to 72 hours on a GridION platform [Oxford Nanopore Technologies] using the MinKNOW software v. 21.05.8 and base called using the super-accurate model (r941_min_sup_g507) with Guppy v. 5.0.11. Downstream consensus sequences were generated using the longread_umi pipeline by Karst et al. 2021^17^ with slight modifications to ensure compatibility with the updated medaka model (r941_min_sup_g507) and the custom barcode sequences. The quality of the consensus sequences was evaluated based on a ZymoBIOMICS Microbial Community DNA Standard [Zymo Research D6306] included together with the samples. With UMI sequence coverage of >= 7x corresponding to Q30+ and >= x14 to Q40+. The exact commands used to generate the consensus sequences was: longread_umi nanopore_pipeline -d input.fq -v 10 -o analysis -s 140 -e 140 -m 1000 -M 2000 -f GGAATCACATCCAAGACTGGCTAG -F AGRGTTYGATYMTGGCTCAG -r AATGATACGGCGACCACCGAGATC -R GACGGGCGGTGWGTRCA -c 3 -p 2 -q r941_min_sup_g507 -t 20 -T 2 -U “3;2;6;0.3”.

### Bacterial and Eukaryotic rRNA gene operon library prep, sequencing and processing

A representative set of 449 samples were selected for both bacterial and eukaryotic rRNA operon sequencing. For bacterial rRNA sequencing PCR was used to amplify ∼4500 bp targeting the 16S and 23S gene using the primers MFD_16S_8F and MFD_23S_2490R. For eukaryotic rRNA operon sequencing PCR was used to amplify ∼4500 bp targeting the 18S and 28S gene using the primers MFD_18S_3NDF and MFD_28S_21R. The PCR reactions contained 10-20 ng DNA input, 1X SuperFi buffer, 0.2 mM dNTPs, 500 nM of each primer, and 2 U of Platinum SuperFi DNA Polymerase [Thermo Fisher Scientific] in a total volume of 50 µL. With the addition of 1X SuperFi GC enhancer when amplifying the eukaryotic rRNA operons.

The PCR-program consisted of initial denaturation at 98 °C for 1 minutes followed by 25 cycles of denaturation (98 °C for 15 seconds), annealing (55 °C for 15 seconds) and extension (72 °C for 3 minutes), followed by a final extension at 72° C for 5 minutes. The PCR products were then purified with SPRI beads [CleanNGS] in a ratio of 0.7x bead/sample. Following 5 minutes of incubation the beads were washed twice in 80 % ethanol and eluted in 20 µL NFW for 5 minutes. The amplicon DNA was barcoded in a second 8-10 cycle PCR reaction using barcoded primers targeting the introduced adapter sequence. The PCR reaction contained 20 ng of the purified amplicon DNA, 1X SuperFi buffer, 0.2 mM dNTPs, 500 nM of forward and reverse primer, and 2 U of Platinum SuperFi DNA Polymerase [Thermo Fisher Scientific] in a total volume of 50 µL. With the addition of 1X SuperFi GC enhancer when amplifying the eukaryotic rRNA operons. The PCR-program consisted of initial denaturation at 98 °C for 1 minutes followed by 8-10 cycles of denaturation (98 °C for 15 seconds), annealing (60 °C for 15 seconds) and extension (72 °C for 3 minutes), followed by a final extension at 72° C for 5 minutes. The PCR products were purified with SPRI beads [CleanNGS] as described above and eluted in 18 µL NFW.

The barcoded amplicons were multiplexed in pools of 92 samples containing a total of 1-2 µg of DNA. The pools were size selected using SPRI ProNex® Chemistry [Promega] with a ratio of 1.2x bead/sample following manufacturer’s protocol and eluted in 125 µL. The size selected and purified pools were shipped for PacBio CCS sequencing on the Sequel II platform [Pacific Biosciences] using the binding kit 3.2. The CCS sequences were further processed using Lima v2.6.0 [Pacific Biosciences] to filter and demultiplex the data. This was done using the hifi-preset ASYMMETRIC and the following settings “--min-score 70, --min-end-score 40, --min-ref-span 0.75, --different, --min-scoring-regions 2”. Furthermore, remaining ligation products were removed by identifying partially remaining adapter sequences after filtering. Subsequently all reads were oriented while removing primer sequences and filtering reads below 3.5 kb or above 6.5 kb using cutadapt^87^ v3.4.

16S rRNA genes corresponding to the V1-V8 region were extracted from the rRNA operons using a custom script (*trim_RNA_operons.sh*) that carry out several steps: First the rRNA operons were truncated to 1450 bp using the usearch^88^ v11 command “fastx_truncate -trunclen 1450”. The trimmed sequences were thereafter trimmed based on the 1392r primer using Cutadapt. Sequences where the primer could not be found were aligned to the global SILVA 138.1 SSURef NR99^18^ alignment using SINA^89^, the aligned sequences were trimmed according to the position of the primer binding sites in the alignment. Truncated sequences were removed using a custom script (*remove_incomplete_seqs_from_sina_aln.py*) that considered sequences that start or end with three or more gaps as truncated. Finally, gaps were removed using the custom script (*Remove_gaps_in_fasta.py*), whereafter the primer- and alignment-trimmed sequences were combined.

Before diversity analysis, 18S rRNA genes, correposing to the position between the 3NDf^90^ and 1510R^91^ primer binding sites, were extracted using a custom script (*trim_euk_RNA_operons_3ndf-1510R.sh*). This script was identical to the script used for processing bacterial rRNA operons except that sequences were trimmed based on the 1510R primer with cutadapt^87^ and the alignment trimmed based on the corresponding position in the SILVA global alignment. The resulting reads were dereplicated using “usearch -fastx_uniques -sizeout” and then resolved into ASVs using “usearch - unoise3 -minsize 2”. An ASV-table was created based on the trimmed reads using “usearch -otutab -zotus”. Taxonomy was assigned to the ASVs using the UTAX version of the PR2 database^26^ v5.0.0, the SINTAX classifier^27^. Phylogenetic diversity of the 18S rRNA genes was determined by clustering the ASVs at 99% identity with “usearch -cluster_smallmem -maxrejects 512 -sortedby other”, followed by mapping against the PR2 database^26^ to determine the percent identity with the closest hit in the database using “usearch -usearch_global -maxrejects 0 -maxaccepts 0 -top_hit_only -id 0 -strand plus”.

### Establishment of the MFD and MFG 16S rRNA gene reference database

The MFG 16S rRNA reference database was assembled from high-quality bacterial and archaeal 16S rRNA genes obtained from several sources: Full-length 16S rRNA gene and rRNA operon amplicons created in this study, SILVA v138.1 SSURef NR99^18^, EMP500^6^, AGP70^17^, MiDAS^2,19^, and Askov/Cologne soil^20^. All bacterial sequences were trimmed between the 8F/27F^92^ and 1391R^93^ primer binding sites and archaeal sequences between the 20F^94^ and the SSU1000ArR^95^ primer binding sites.

High-quality bacterial and archaeal sequences were obtained from the SILVA 138.1 SSURef NR99^18^ ARB-database by exporting them separately in the “fastawide” format after terminal trimming between position 1,044 and 41,788 (Bacteria) and 1,041 and 32,818 (Archaea) in the global SILVA alignment, corresponding to the primer binding sites. A custom script (*Extract_full-length_16S_rRNA_genes_from_SILVA_alignments.sh*) was used to remove truncated sequences based on the presence of terminal gaps in the exported FASTA-alignments. Finally, sequences that contained N’s were removed and U’s replaced with T’s using two custom scripts (*Remove_seqs_with_Ns.py* and *replace_U_with_T.py*).

Due to the large number of trimmed 16S rRNA gene reads, ASVs were resolved for each data set individually. Sequences were dereplicated using “usearch -fastx_uniques -sizeout” and then resolved into ASVs using “usearch -unoise3 -minsize 2”. ASVs from all individual datasets were combined together with the 16S sequences from SILVA and dereplicated using “usearch - fastx_uniques”. The ASVs were sorted based on their abundance across the MFD dataset by mapping the trimmed sequences against the ASVs with “usearch -search_exact -dbmask none - strand plus -matched”. The matched sequences were dereplicated using “usearch -fastx_unique - sizeout”, and sequences which did not originate from the MFD samples were appended. The complete database was processed using AutoTax^21^ v1.7.6 to create the MFG database. Because the MFG database was found to contain sequences representing chloroplast and mitochondrial 16S rRNA genes as well as pseudogenes, we provided additional filtering to create the final MFG database. These included removing i) all sequences that shared less than 70% identity with their closest match in the SILVA 138.1 SSURef NR99 database^18^ ii) ASV with de novo placeholder names and best hits in SILVA against Mitochondria or Chloroplast; iii) ASV with de novo phyla placeholder names and top hits against Rickettsiales and less than 75% identify; iv) ASVs representing de novo phyla covered by only a single ASV; v) ASVs with a better hit in the MIDORI2 mitochondrial database^96^ then in SILVA, which at the same time share >75% identity (>1000bp alignment length) to the MIDORI2 hit. Finally, ASVs which were assigned to de novo phyla but shared >70 % identity with a Patescibacteria hit were assigned to Patescibacteria.

The MFD database was obtained by subsetting the MFG one based on samples only found among the MFD datasets. A 98.7% identity clustered version of the MFG database was obtained directly from the AutoTax output (temp/SILVA_FLASV-S_centroids.fa).

### Database evaluation

To evaluate the coverage of the MFG database, we mapped 16S gene fragment reads extracted from the 10km EU reference grid of Denmark representative metagenomes and 16S rRNA gene V4 OTUs clustered at 99% identity from the Global Prokaryotic Census project^13^ against our databases as well as SILVA 138.1 SSURef NR99^18^, GreenGenes2^28^, and the complete 16S rRNA database from GTDB v220^29^ using “usearch -usearch_global -maxaccepts 512 -maxrejects 0 -top_hit_only -id 0”. The mapping data was used to calculate the percentage of reads with hits above specified identity thresholds representing the different taxonomic ranks^23^. The reads were also classified using the different databases and the SINTAX classifier^27^, and the percent of reads classified at different taxonomic ranks was determined. The analysis was conducted using R^77^ v4.3.2 using tidyverse^97^ v2.0.0 and vegan^98^ v2.6-6.1.

### Short-read 16S rRNA gene classification

Hidden markov models (HMM) were made from Rfam^99^ v14.7 seed alignments for Archaea (RF01959), Bacteria (RF00177) and Eukarya (RF01960) using hmmbuild (HMMER^100^ v3.3.2). Metagenomic reads from rRNA gene fragments were extracted from the quality-filtered metagenic reads using the constructed models with nhmmer^101^ (HMMER^100^ v3.3.2) with settings “--incE 1e-05 -E 1e-05 --noali”. In the case of multiple hits for the same metagenomic read, the best domain hit was selected based on the bit-score. The 16S reads were filtered for hits within the region between position 27F^92^ and 1391R^93^ primer binding sites for Bacteria and the 20F^94^ and SSU1000ArR^95^ primer binding sites for Archaea. The 16S reads were taxonomically annotated using the SINTAX classifier^27^ and the MFG database with the confidence cutoff set to 0.8. The output was aggregated to the individual taxonomic levels using R^77^ v4.4.1 and the tidyverse^97^ v2.0.0 package. Reads not classified at the given taxonomic level were assigned to “Unclassified”.

### Diversity calculations

Using usearch^88^ v11 all V1-V8 MFD 16S gene reads were mapped against the species-level representative OTUs of the MFG database at 98.7% identity threshold and downstream analysis done using R^77^ v4.4.1. The alpha diversity was only calculated from samples with more than 9,000 reads, which were randomly subsampled without replacement using vegan^98^ v2.6-6-1 (function *rrarefy*) to a depth of 9,628 reads. The gamma diversity was estimated after transformation to incidence data using rarefaction and extrapolation with Hill numbers as implemented in the iNEXT^102^ v3.0.1 package (function *iNEXT*). The species richness and the richness of common species (Shannon diversity, i.e., the exponential of the Shannon index) was estimated using order *q* = 0 and *q* = 1 respectively. The gamma diversity was estimated for the total set of samples (Denmark) and habitat specific sets (MFDO1). For the total set the endpoint of extrapolation was set to twice the set size and for habitat specific estimates the endpoint was fixed at 100 samples.

Principal Coordinate Analysis (PCoA) was performed on the 16S gene fragments aggregated on the genus level from samples with more than 1,000 reads, i.e., 9,812 samples and subjected to random subsampling without replacement using ampvis2^103^ v2.8.9 (functions *amp_load* and *amp_subset_samples*). Abundances were Hellinger-transformed using vegan^98^ v2.6-6.1 (function *decostand*) before calculation of Bray-Curtis distance calculated with the package parallelDist^104^ v0.2.6 (function *parDist*) before performing the principal coordinate decomposition with the ape^105^ v5.8 package (function *pcoa*). In order to investigate the effect of spatial autocorrelation on community composition we performed a distance decay analysis across the levels of the MFD ontology. The autocorrelation was modelled using a logarithmic decay model of Hellinger-transformed distance as a function of spatial distance calculated with the codep^106^ v1.2-3 package (function *gcd.hf*). The modelling was restricted to samples with reliable coordinates and MFDO1 habitats showing spatial separation of samples, with the exclusion of Bornholm and limited to spatial distances <300 km. A pseudo-count of 0.1 m was added to samples with zero spatial distance. In order to address the effect of spatial autocorrelation on community composition the data was spatially thinned using the 10-km reference grid of Denmark^81^, by selecting a sample representative of each MFDO1 habitat having the lowest Bray-Curtis distance to other samples of that habitat in the same cell. This led to a spatially thinned subset of 2,122 samples. From this subset beta diversity was visualised using the stats^77^ v4.4.1 package (function *hclust*, method ward.D2) on the between-group Bray-Curtis distances, with the confidence of the group-splits was calculated using 100 iterations with the bootstrap^107^ v0.1 package (function *bootstrap*).

### Habitat classification

The habitat classification analysis was performed using R^77^ v4.2.3. The microbial relative abundances at the genus level of the 16S gene fragments were summarised at higher taxonomic levels (family to phylum) by summing up the abundances of populations with the same taxonomy. Taxa observed with a relative abundance >0 in at least 25 samples were retained for subsequent analysis. The resulting five tables were screened for multicollinearity using the function *vifcor* (th=0.7) from the package sdm^108^ v1.1_18 wrapped in a block-wise script for efficiency. Considering the block-wise implementation it is not guaranteed to find the optimal solution, but the shuffling at each iteration increases the chances of approaching it. The ontology was used to create five different target variables, one for each ontology level, but only classes with at least 50 observations and whose class names were not ending with “NA” were retained for modelling. The modelling was carried out with the package tidymodels^109^ v1.1.1, which allows the creation of model workflows. The functions cited in this section belong to tidymodels^109^ unless otherwise stated. The predictor variables (microbial relative abundances) were centred on the mean, scaled to unit variance and used to build a random forest model (from the package ranger^110^ v0.16.0, using default parameters). The model was validated 5 times with a 5-fold cross validation using the function *vfold_cv* with v = 5 and repeats = 5; where the evaluation metrics were collected on the 5th split of the data, on rotation, leading to 25 evaluation values per model. The individual fit information was extracted with the *extract_fit_parsnip* function workflow and the variable importance computed with the command *vi* (using default parameters) from the package vip^111^ v0.4.1. Global metrics such as F1 (micro and macro), PR AUC and Kappa were collected after the cross validation using the function *collect_metrics*. We ran a total of 25 combinations between predictors (phylum, class, order, family, genus), and targets (Sample type, Area type, MFDO1, MFDO2, MFDO3) with the function *fit_resample* and specifying the above mentioned cross validation, evaluation metrics and fit extraction as resample, metrics and control parameters, respectively. The analysis of the false negatives was performed by collecting, for each class in each fold and iteration the number of associated misclassified samples, using the function *conf_mat* from the package yardstick^112^ v1.3.0. A null distribution of false negatives was computed by multiplying the previous number by the fraction of samples in each class (except for the class in exam). A two-tailed paired t-test was performed using the function t_test from the package rstatix^113^ v0.7.2.

### Core analysis

Abundant and prevalent genera were identified from the spatially thinned dataset across the MFD ontology using a prevalence and mean relative abundance filter of 50% and 0.1% respectively. Investigation of shared genera was performed using UpSetR^114^ v1.4.0 and ComplexUpset^115^ v1.3.3.

### Metagenomic assembly and binning

Trimmed shallow metagenomic reads were assembled using MegaHit^116^ v1.2.9 with the following options: “--k-list 27,43,71,99,127 --min-contig-len 1000”. Metagenomes smaller than 1 Mbp were omitted from further processing. In total, 10,042 assemblies were used for genome recovery.

To maximise the recovery of low coverage MAGs, SingleM^42^ v1.0.0beta7 “pipe” subcommand with the default GTDB R214 metapackage^117^ was first run on the metagenomes to generate archive OTU tables of single-copy marker gene sequences (combined with SingleM summarise across all samples). Bin Chicken^118^ v0.9.6 was then run on those tables using the command “ibis coassemble –max-coassembly-samples 1 –max-recovery-samples 10 –singlem-metapackage S3.2.1.GTDB_r214.metapackage_20231006.smpkg.zb”, to match the single copy protein sequences across samples and choose the 10 most similar samples for each sample for multi-sample binning. The reads for the selected samples were mapped to the assemblies using Minimap2^119^ v2.24 with the “-ax sr” option and SAMtools^120^ v1.16.1 with “samtools view -Sb -F 2308 - | samtools sort” options. The “jgi_summarize_bam_contig_depths” command of MetaBAT2^121^ v2.12.1 was used on the mapping files to acquire contig coverage values, which were used as input for binning via MetaBAT2^121^ with the following options: “-m 1500 -s 100000”.

The recovered MAGs were quality-assessed using CheckM2^122^ v1.0.2 to acquire MAG completeness, contamination values and were taxonomically classified using GTDB-Tk^123^ v2.3.0. Bakta^124^ v1.8.1 with Bakta database v5.0 (type “full”) was used to annotate the MAGs and acquire bacterial rRNA and tRNA counts, while for archaeal MAGs, tRNAscan-SE^125^ v2.0.9 and barrnap^126^ v0.9 with the corresponding archaeal databases were used to acquire tRNA and rRNA counts. CheckM2^122^ completeness and contamination values, together with the observed rRNA and tRNA counts, were used to classify the MAGs according to MIMAG guidelines^22^ and only medium or high quality MAGs were kept for further analysis. The MAGs were de-replicated using dRep^127^ v2.6.2 with “-sa 0.95 -nc 0.4” settings. MAG coverage values were calculated using CoverM^128^ v0.6.1, with the “-m mean” setting.

Representation of the metagenomic microbial community by these short-read-assembled MAGs in each sample was assessed using SingleM^42^ v0.16.0 (metapackage used was supplemented with MAGs beyond GTDB https://doi.org/10.5281/zenodo.10360137) through the “appraise” subcommand that compares the OTUs of 59 single-marker copy genes found in the metagenomic reads and MAGs. Briefly, SingleM “pipe” subcommand was first applied to individual trimmed short-read metagenomes (with a minimum of 100 bp) and individual short-read assembled MAGs to identify OTU sequences of these 59 marker genes in both types of datasets. For each type of dataset (i.e., short-read data and MAGs), OTU tables were then combined using SingleM “summarise” and passed through SingleM “appraise” subcommand (under the default cutoff for species level estimates) to compare the OTU tables of both data types for determining the bacterial and archaeal community recovered by the short-read assembled MAGs.

### Novelty and prokaryotic fraction estimation

To assess sample novelty and prokaryotic fraction in each shallow metagenome, microbial profiles were estimated by SingleM^42^ v0.16.0 (metapackage: supplemented with MAGs beyond GTDB https://doi.org/10.5281/zenodo.10360137) using the SingleM “pipe” subcommand. Essentially, OTUs of 59 single-copy marker genes were first identified from the shallow metagenomes and compared against those found in the reference genome database to assign taxonomic classifications for the identified OTUs. The overall taxonomic profile for each metagenome was then condensed from the OTU tables of all the different markers. Sample novelty was expressed in terms of the proportion of microbial community characterised by the current reference genome database that encompassed most recent advances in genome mining. This was achieved by taking the ratio between the sum of the total coverage at the species level and the sum of total coverage at all taxonomic levels based on the condensed taxonomic profiles (referred to “known species fraction” thereafter)^42^. To compare novelty between metagenomes from MFD and NCBI public metagenomes, published SingleM profiling results for NCBI datasets^42^ were used for comparison with the MFD datasets. NCBI datasets with metadata habitat labelling in the original paper of “marine”, “freshwater”, “aquatic”, “drinking water”, “freshwater”, “groundwater”, “lake water”, “wastewater” groups were grouped in into NCBI_Water, and NCBI datasets of “sediment”, “marine sediment”, “freshwater sediment” groups were grouped into NCBI_Sediment. To approximate the prokaryotic fraction of a sample, the proportions of bacterial and archaeal reads present in a metagenome estimated using the SingleM “microbial_fraction” subcommand were added up.

To investigate the improved species representation of microbial community by the short-read constructed MAGs, these MAGs, together with MAGs from beyond GTDB^10^ (SPIRE^8^, SMAG^11^, GEM^7^, UHGG^9^, ocean MAGs^12^) were added to the default GTDB R214 SingleM metapackage (https://doi.org/10.5281/zenodo.8419620). This was done using the SingleM “supplement” subcommand applying dereplication at 95% similarity and CheckM2 quality filtering of 50% completeness and 10% contamination for all MAGs to be supplemented. In total, 4,109/19,253 dereplicated novel species representatives from MFD MAGs were added to the new metapackage. Microbial community was then re-analyzed based on this new metapackage supplemented with the short-read constructed MAGs using the SingleM “renew” subcommand.

### Gene-centric investigation of short-read metagenomes (database from GEM and GTDB, curation and HMM generation for GraftM)

A dereplicated set of the species representatives from the genome taxonomy database^10^ release 214 and the genomic catalogue of Earth’s microbiomes^7^ was created using fastANI^129^ v1.32 removing all GEMOTU genomes with >96 % average nucleotide identity over >50 % aligned fragments to a GTDB species representative. The resulting 97,227 genomes (85,205 GTDB and 12,022 GEMOTU) were annotated using anvi’o^130^ v8 with gene calling using Prodigal^131^ v2.63.

The total protein complement of the 97,227 genomes was used as a query in a DIAMOND^132^ v2.1.8 search against preliminary protein datasets of *N*arG/NxrA and *A*moA/PmoA with a score cutoff of 100. True positive hits were selected using an alignment score ratio approach as previously described^133,134^, resulting in 1068 (AmoA/PmoA) and 11,500 (NxrA) sequences.

For further analyses, the AmoA/PmoA sequences were split in 823 bacterial and 245 archaeal sequences and the NarG/NxrA sequences were split into a cytoplasmic group (NarG, 10,254 sequences) and two predicted periplasmic groups (*Nitrospira* NxrA-like, 665 sequences and *Nitrotoga* NxrA-like, 581 sequences) based on phylogeny.

Multiple sequence alignment of protein sequences was performed with MAFFT^135^ v7.490. TrimA^l136^ v1.4.1 was used to trim minimum 20% aa representation. IQ-TREE ^137^ v2.2.0.3 was used to generate phylogenetic trees of protein sequences, utilising the ultrafast bootstrap approximation option and 1000 iterations. For NxrA/NarG and bacterial AmoA trees, the ModelFinder option was applied (Best-fit models pNXR: LG+R6, cNXR: LG+F+R10, Bacterial amoA: LG+F+R8). The model applied for the archaeal AmoA tree was WAG+R. ARB^138^ v7.0 was used to re-root and group trees, followed by visualisation in iTOL ^139^ v6. Inclusion of protein sequences of NxrA and AmoA obtained through this study was filtered based on length of the protein product, with a cutoff of 200aa for AmoA and 600aa for NxrA. Gene phylogeny was ascertained using GraftM^140^ v0.14 on the forward shallow metagenomic reads with search mode “hmmsearch+diamond” along with a conditional E-value threshold of 1e-10. The output was aggregated to the individual clades using R^77^ v4.4.1 and the tidyverse^97^ v2.0.0 package. Read abundance was normalised by HMM-alignment length (Bacterial AmoA: 744nt, Archaeal AmoA: 648nt, pNXR: 3411nt, cNXR: 4092nt) and sequencing depth of prokaryotic reads determined by SingleM, reported in reads per kilobase million (RPKM). Samples were filtered to include only samples from MFDO1 habitats “Sediment, Natural, Freshwater”, “Sediment, Urban, Freshwater”, “Soil, Agriculture, Fields”, “Soil, Natural, Bogs, mires and fens”, “Soil, Natural, Forests”, “Soil, Natural, Grassland formations”, “Soil, Urban, Greenspaces”, and “Water, Urban, Wastewater”. Read abundances were Hellinger-transformed using vegan^98^ v2.6-6.1 (function *decostand*) before calculation of Bray-Curtis distance with the package parallelDist^104^ v0.2.6 (function *parDist*) and performing the principal coordinate decomposition with the ape^105^ v5.8 package (function *pcoa*). Samples were clustered within MFDO2 habitats using hierarchical clustering with the stats^77^ v4.4.1 package (function *hclust,* method ward.D2).

### Phylogenomic investigation of Nitrifier genera, and Nitrobacter-like NxrA containing MAGs compared to NxrA gene phylogeny

*Xanthobacteraceae* family MAGs that had been identified harbouring *Nitrobacter*-like NxrA sequences of at least 600aa in length were selected for phylogenetic analysis, alongside species representatives of the SR MAGs that met >90% genome completeness and <5% genome contamination thresholds using CheckM2^122^ v1.0.2 and belonged to the genus level GTDB groups BOG-931, *Bradyrhizobium*, JAFAXD01, *Pseudolabrys* and VAZQ01. The single recovered *Nitrobacter* MAG from this study was also included, as well as representatives from GTDB for context, for example the species representatives present for BOG-931, JAFAXD01, VAZQ01 and manually selected isolates from *Bradyrhizobium* and *Pseudolabrys*. The specific Nxr groups were assigned based on the updated GraftM package and classification tree. The phylogenomic tree was constructed using GTDB-Tk^123^ v2.3.2 R214 and the de_novo_wf on all bacterial MAGs. The multiple sequence alignments of 120 single copy proteins were subset to the genomes of interest using fxtract^141^ v2.3. This alignment was used as input for IQ-TREE^137^ v2.1.2 using the WAG+G model and -B 1000 using UFBoot. The tree was visualised in ARB^138^ v7.0 for rerooting by the *Methylopilaceaea* isolate outgroups and analysis, and further processed in ITOL^139^ v6.

Phylogenomic trees were created by subsetting the GTDB-Tk de_novo_wf trees for the bacteria and archaea for known nitrifier genera as well as the putative genera examined in this study. Trees were investigated and refined using ARB and ITOL as above, with final refinements in Inkscape.

The cytoplasmic NxrA tree was built using *Nitrobacter*-like NxrA sequences of at least 600aa in length from *Xanthobacteraceae* family MAGs, along with known NxrA from *Nitrococcus* and *Nitrobacter*, and outgroup NxrA sequences from *Nitrospira*, *Scalindua*, and *Brocadia*. Alignment, trimming and tree-generation was done using MAFFT^135^ v7.490, TrimA^l136^ v1.4.1, IQ-TREE^137^ v2.2.0.3 with ultrafast bootstrap approximation and 1000 iterations, using substitution model LG+F+R10.

### Metabolic investigation of nitrifier MAGs

Metabolic reconstruction of the short-read MAGs was conducted using DRAM^142^ v1.4.6 and the KEGG database^143^ release 109. ‘DRAM.py annotate’ was run using the default settings, followed by ‘DRAM.py’ distil. Key nitrification and CBB cycle genes were searched based on KO identifiers in annotations.tsv file and incorporated into the genome trees. Gene-synteny was displayed using R^77^ v4.4.1 and the gggenes^144^ v0.5.1 package.

### Other R-packages

Other R-packages used in the conducted analysis not listed elsewhere are readx^l145^ v1.4.3, data.table^146^ v1.15.4, wesanderson^147^ v0.3.7, viridisLite^148^ v0.4.2, ggpubr^149^ v0.6.0, patchwork^150^ v1.2.0, ggpmisc^151^ v0.5.6 and hexbin^152^ v1.28.3.

