## Supplementary Information for "Microflora Danica: the atlas of Danish environmental microbiomes"

**Affiliations**

Mads Albertsen

Center for Microbial Communities, Department of Chemistry and Bioscience,

Aalborg University, Fredrik Bajers Vej 7H, 9220 Aalborg, Denmark

Per Halkjær Nielsen

Center for Microbial Communities, Department of Chemistry and Bioscience,

Aalborg University, Fredrik Bajers Vej 7H, 9220 Aalborg, Denmark;

**Table of content**

Supplementary Figure 1-8

Supplementary Table 1-4

Supplementary Note 1-6

Supplementary References

### Supplementary Figures

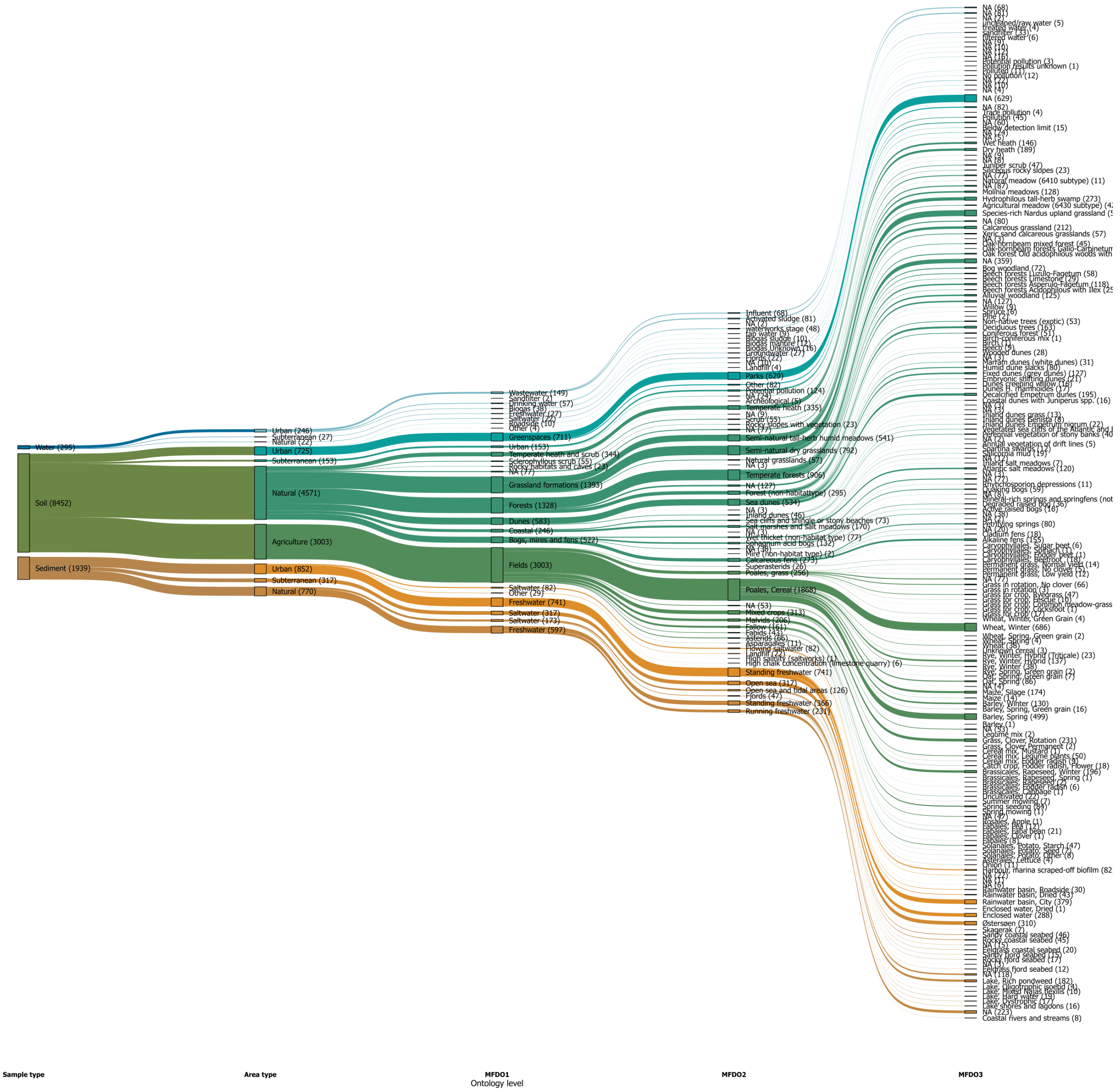


**Supplementary Figure 1: Sample repartition in the five levels of the habitat ontology.** Full version of **Figure 1c**, depicting the full structure of the habitat ontology. The colours of the branches represent the node of origin for the first two levels (i.e., first level of branches = Sample type, second level of branches = Sample type + Area type). The following two levels of branches are coloured as Sample type + Area type. Each habitat category is followed by the number of samples for that category in parentheses.

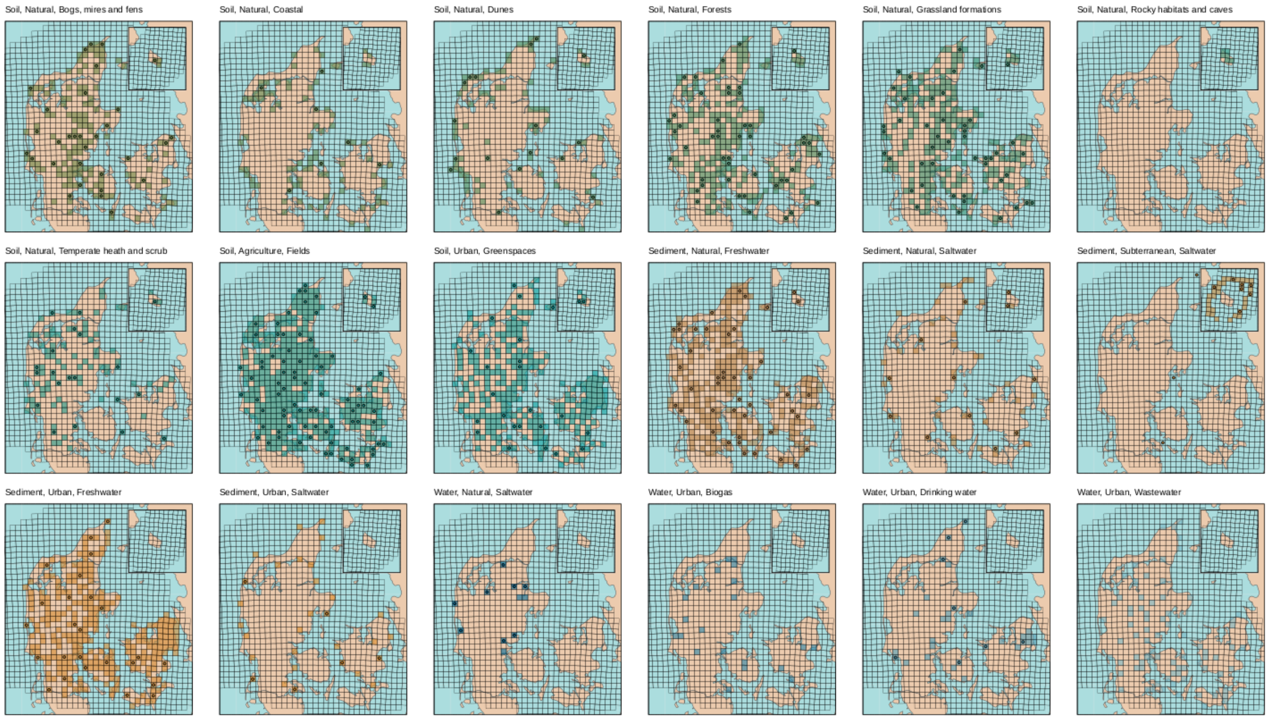


**Supplementary Figure 2: Distribution of the samples per MFDO1 habitats in the 10km EU reference grid of Denmark.** The map of Denmark follows the same criteria and scales as **Figure 1a**, including the cutout for the Bornholm island. Each facet includes only samples of the relative MFDO1 category as indicated in each title and the colouring follows the scheme in **Figure 3b**. A total of 18 (9,925 samples) of MFDO1 habitats showed a broad geographical distribution of samples across the country. An exception for one of the 18 is MFDO1 "Rocky habitats and caves", with samples only found on the island of Bornholm, which is geologically different from the rest of Denmark. The 10 km EU reference grid of Denmark[81] was overlaid to the map and each cell where at least a metagenomic sample was present has been shaded. Cells including at least a FL16 sample have been marked with a dot in the centre.

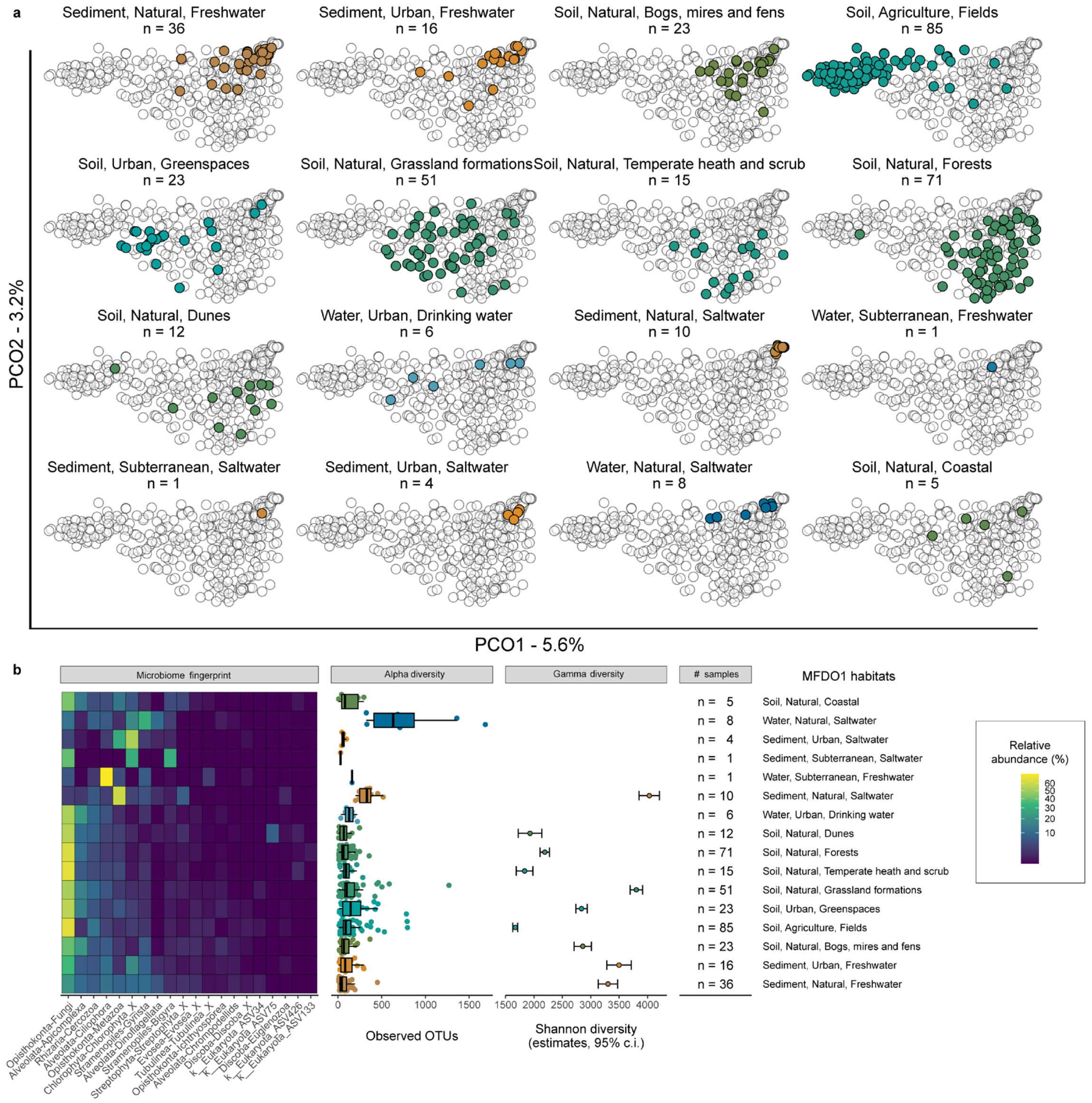


**Supplementary Figure 3: Eukaryotic diversity of the Danish habitats. a.** Ordination of the 18S rRNA dataset. 18S rRNA sequences from rRNA operon sequencing of 449 representative samples are arranged using PCoA and 16 selected MFDO1 habitats are highlighted in different panels and colours. The visualisation depicts the first two components, accounting for 5.6% and 3.2% of explained variance, respectively. **b.** Diversity overview of the habitats. The selected 16 MFDO1 habitats are represented in the rows of the multi-facet plot, a heatmap shows the relative abundance of the 20 most abundant Divisions-Subdivisions, taxonomy is inferred with the PR2 database[26]. Each facet addresses a different aspect of diversity.

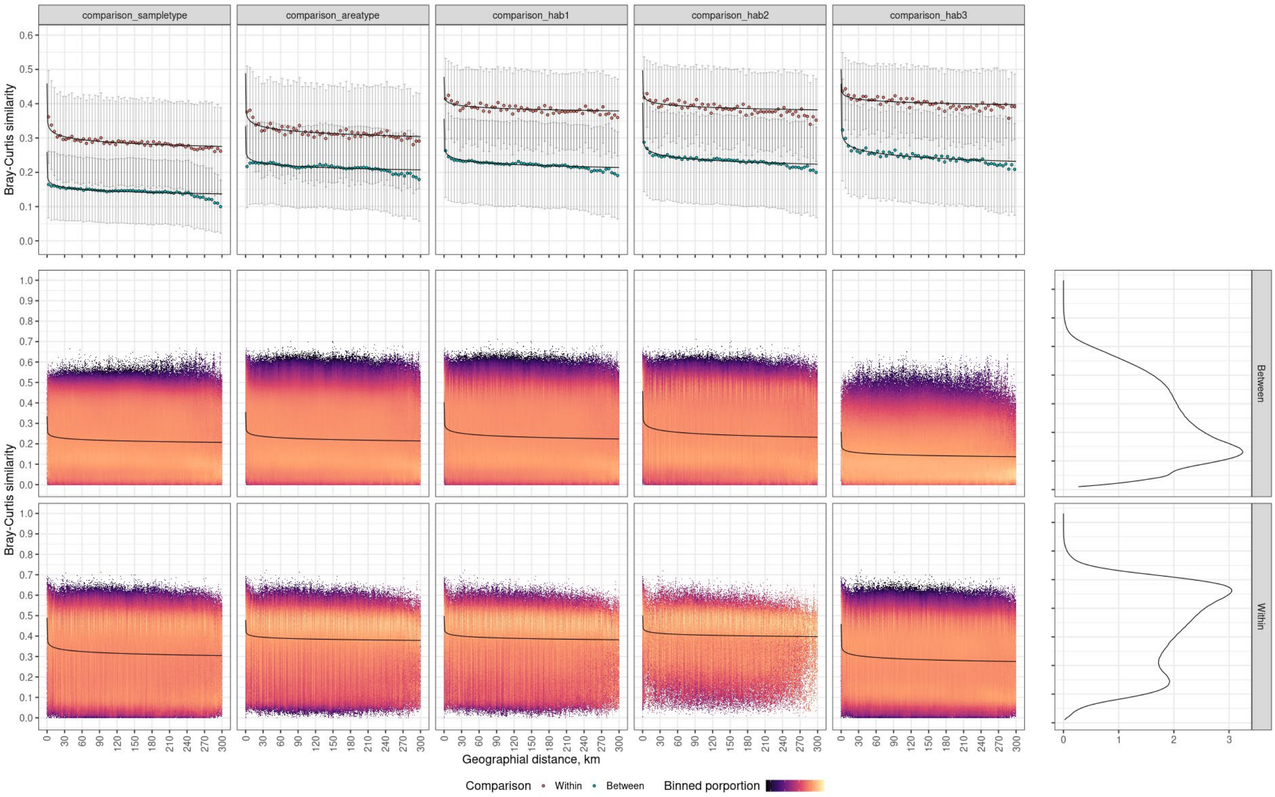


**Supplementary Figure 4: Distance decay analysis.** The analysis was conducted on genus-aggregated 16S rRNA data subjected to random subsampling without replacement (1,506) reads. The samples were filtered to samples with information on MFDO3, reliable coordinates (except from masked Agricultural), exclusion of samples from Bornholm and limited to distances below 300 km. Samples were further filtered for MFDO3 habitats with more than 9 representatives. This left 9,121 samples (6,325,297 reads) for the calculation of Hellinger-transformed Bray-Curtis and geographical distances (a total of 371,917,507 comparisons). The influence of spatial distance on the microbial community in the metagenome samples was negligible, except at short distances (<10km).

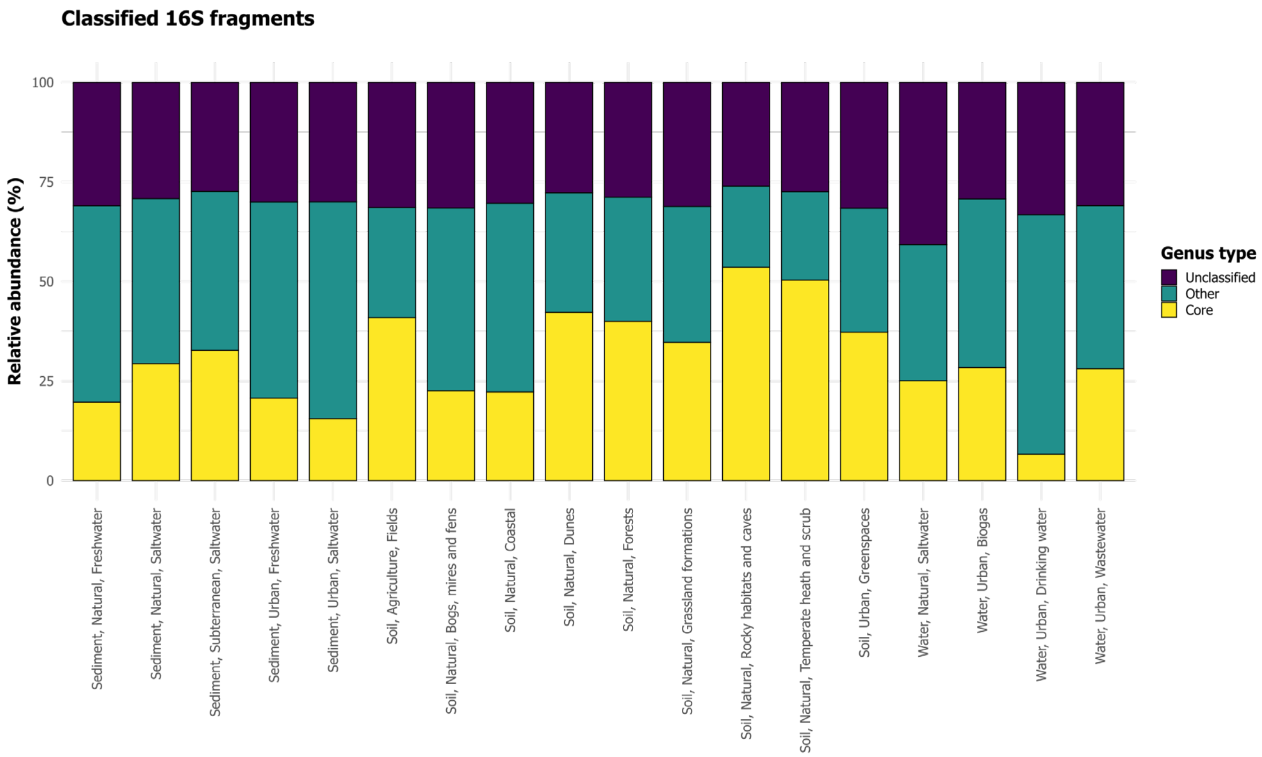


**Supplementary Figure 5. Relative abundance fraction of core organisms.** Stacked bar plot of cumulative relative abundance of the identified core and non-core genera as well as the reads unclassified at the genus level across MFDO1 habitats.

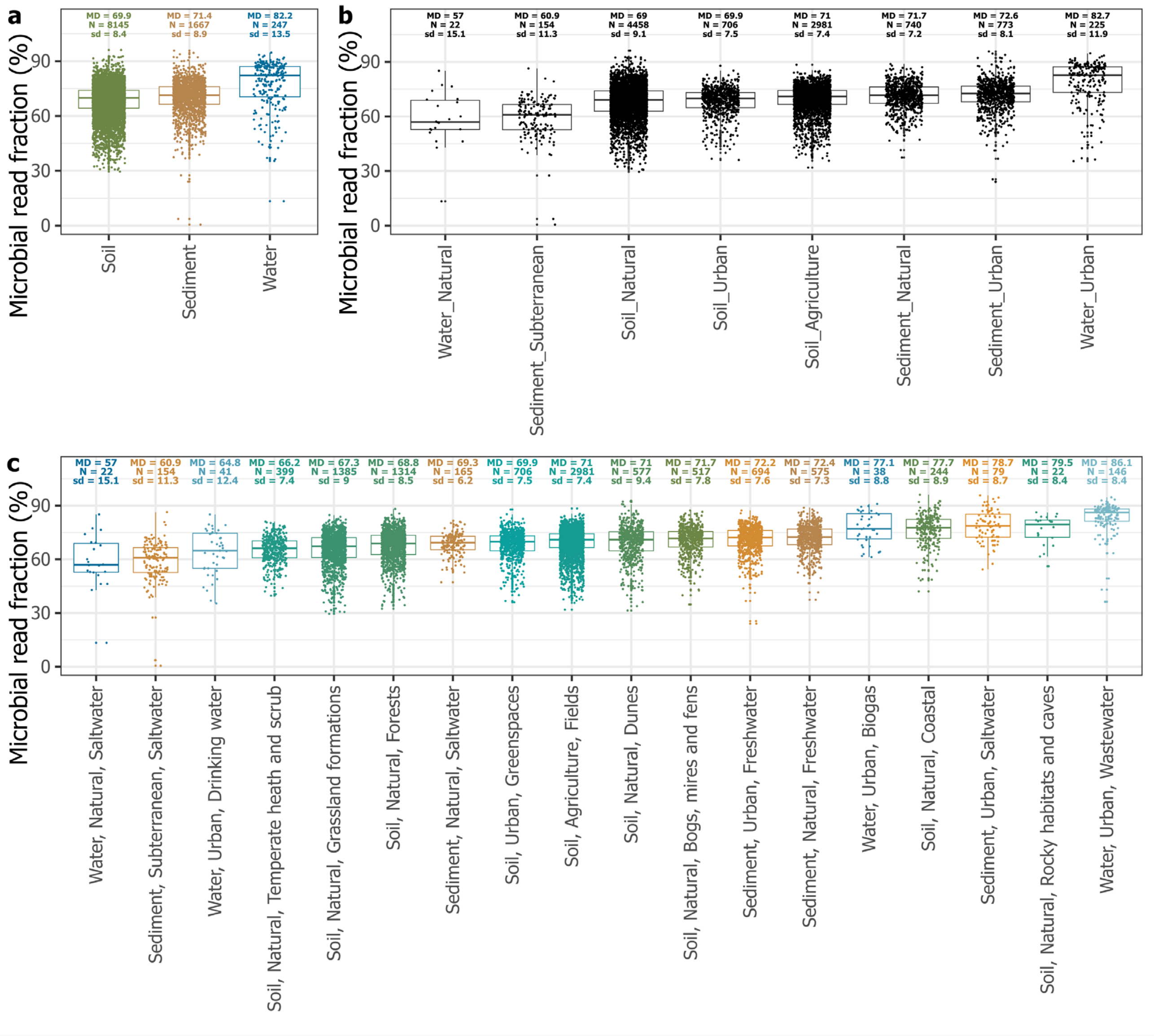


**Supplementary Figure 6. Boxplot of the proportions of prokaryotic (bacterial and archaeal) reads** across different **a.** sample types; **b.** area types; and **c.** habitats. Asterisks and “ns” indicate statistical significance of differences between groups by Mann-Whitney U test with Bonferroni-adjusted $p$-values: ns: $p > 0.05$, *: $p \leq 0.05$, **: $p \leq 0.01$, ***: $p \leq 0.001$, and ****: $p \leq 0.0001$. MD: median, N: sample number, sd: standard deviation.

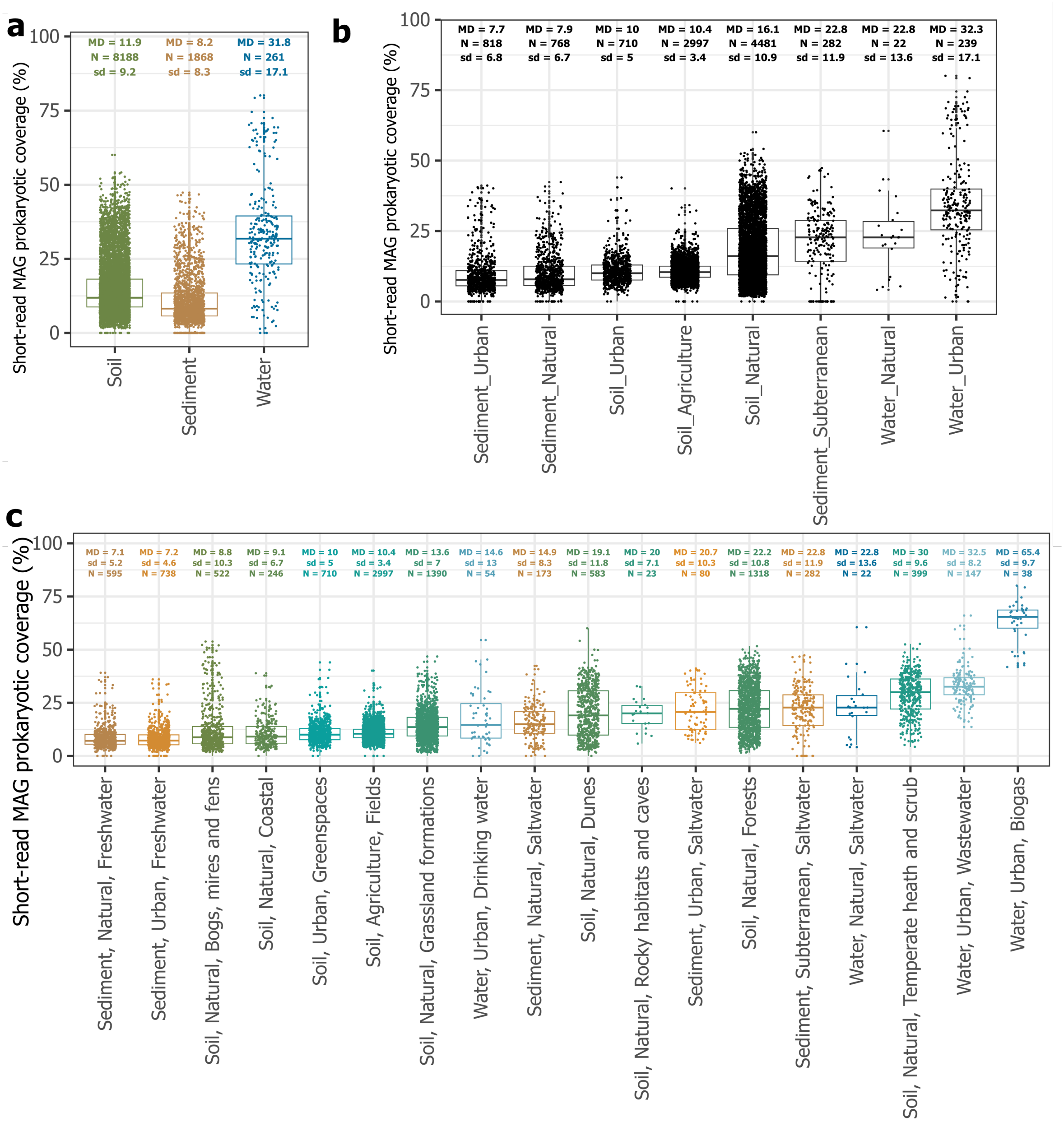


**Supplementary Figure 7. Boxplot of the proportions of prokaryotic community coverage by MAGs reconstructed from short-read metagenomes** across different **a.** sample types; **b.** area types; and **c.** habitats. MD: median, N: sample number, sd: standard deviation.

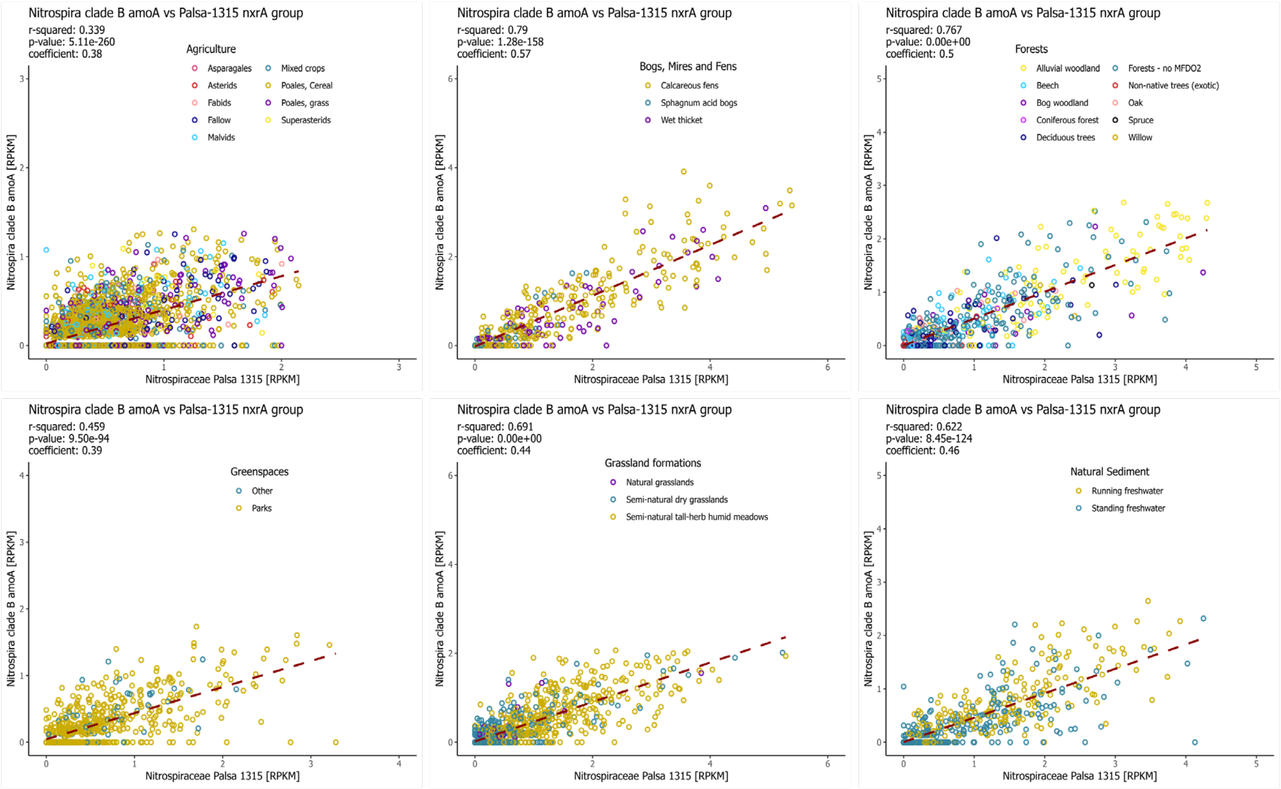


**Supplementary Figure 8. Correlation of gene read abundance of comammox clade B *amoA* and Palsa-1315 *nxrA*.** Relative gene read abundance in selected MFDO1 habitats ‘Agriculture’, ‘Bogs, mires and fens’, ‘Forests’, ‘Urban Greenspaces’, ‘Grassland formations’, and ‘Natural Sediment’. Read abundance is reported in reads per kilobase million [RPKM], and normalised to the fraction of microbial reads determined by SingleM. A least squared distance linear model was used to calculate the $R^2$, correlation coefficient, and p-value. Samples with RPKM > m + 3σ within each MFDO1 have been removed.

### Supplementary Tables

**Supplementary Table 1. Number of core genera across the MFD ontology.** The number of core genera identified across the five levels of the MFD ontology. Columns to the right of an ontology level refers to the genera identified for that ontology level habitat categories.

**Supplementary Table 2. Prevalence of core genera across the MFD ontology.** List of core genera identified in the MFDO3 habitats. Prevalence corresponds to the number of different MFDO3 habitats each genera was identified in. The relative abundance of each genera is reported as median, mean and standard deviation of mean MFDO3 habitat specific relative abundance.

**Supplementary Table 3. Table summarising the fraction of the community (%) assigned to the species level across different sample types in MFD and NCBI datasets.** SingleM metapackage used was the version inclusive of MAGs beyond GTDB (DOI 10.5281/zenodo.10360136). Sample_N: sample number, Std. Dev.: standard deviation, RSD: relative standard deviation. NCBI datasets of "marine", "freshwater", "aquatic", "drinking water", "freshwater", "groundwater", "lake water", "wastewater" groups were grouped in into NCBI_Water, and NCBI datasets of "sediment", "marine sediment", "freshwater sediment" groups were grouped into NCBI_Sediment, following singleM paper NCBI dataset metadata.

**Supplementary Table 4. Table summarising the fraction of the community (%) assigned to the species level across different sample types in MFD datasets with and without the inclusion of MFD MAGs.** For the inclusion of MFD MAGs, SingleM metapackage containing MAGs beyond GTDB (DOI 10.5281/zenodo.10360136) was supplemented with MFD MAGs. SingleM metapackage containing MAGs beyond GTDB (DOI 10.5281/zenodo.10360136) without supplement of MFD MAGs was used as the baseline comparison point for assessing the novelty of MFD metagenomes without MFD MAGs.

### Supplementary Notes

#### Supplementary Note 1: Taxonomic thresholds for 18S rRNA genes

In 2014, Yarza[23] et al proposed 16S rRNA gene-based taxonomic thresholds for bacteria and archaea. Although not considering different evolutionary rates across the tree of life, these thresholds provided valuable insight into the prokaryotic diversity, and provided a framework for assigning provisional taxonomy at higher ranks to uncultured bacteria and archaea. Unfortunately, such thresholds do not exist for eukaryotic 18S rRNA genes but would be of similar value. To determine statistically supported thresholds of eukaryotic 18S rRNA genes, we applied a similar approach as originally used by Yarza et al.[23]. 18S rRNA gene sequences and their corresponding taxonomy was downloaded from the PR2 database[26] v5.0.0. The sequences were trimmed corresponding to the region between the 3NDf[90] and 1510R[91] primer binding sites, and truncated sequences removed, using the same pipeline as we used to extract 18S rRNA genes from our eukaryotic rRNA operons. Next, we performed an all against all mapping of the sequences with usearch -usearch_global -maxaccept 0 -maxrejects 0 -top_hit_only. The mapping data was used to determine, for each sequence, the maximum sequence identity to another sequence within each rank and across ranks. This data was hereafter summarised to provide the median values and and the interval of 90% most common values for all sequences (**Supplementary Note Table 1**). Based on these data we proposed taxonomic thresholds for species, genera, and families, considering both the with and across rank diversity. No reliable threshold could be determined for ranks above the family level.

**Supplementary Note Table 1. Taxonomic thresholds of Eukaryotes.** Results based on Eukaryotic 18S rRNA genes (3NDf[90] to 1510R[91]) obtained from the PR2[26] database v5.0.0. Values are given as the median and the interval of 90% most common values (in parentheses) of highest identity within and across the specified ranks.

| Taxonomic ranks | Number of reference sequences | Maximum sequence identity within the rank | Maximum sequence identity across the ranks | Thresholds for assignment and clustering |
|---|---|---|---|---|
| **Species** | 17380 | 100.0 (97.8-100.0) | 99.3 (89.8-100.0) | 99 |
| **Genus** | 26858 | 99.9 (95.9-100.0) | 98.2 (86.1-100.0) | 97 |
| **Family** | 27477 | 99.9 (93.5-100.0) | 93.1 (79.6-99.5) | 93 |
| **Order** | 25966 | 99.8 (93.1-100.0) | 91.9 (78.9-99.1) | NA |
| **Class** | 31555 | 99.8 (93.1-100.0) | 90.8 (78.7-98.8) | NA |
| **Subdivision** | 22922 | 99.8 (92.4-100.0) | 87.9 (72.2-97.8) | NA |
| **Division** | 27754 | 99.8 (93.0-100.0) | 88.4 (70.8-98.0) | NA |
| **Supergroup** | 27647 | 99.8 (93.0-100.0) | 88.0 (71.0-98.0) | NA |

### Supplementary Note 2: Bacterial diversity

As land-use perturbation, i.e. land management, generally leads to the reduction of above-ground biodiversity[32], we wanted to investigate whether the same could be observed for the microbial communities across the soil habitats. The investigated soil habitats span a gradient of land-use perturbation, which has also been used to investigate continental and global microbial diversity, which found lower bacterial and fungal diversity in less-disturbed environments (woodlands) compared to grasslands and highly-disturbed environments (croplands)[31,30].

Besides these habitats our study included two less-disturbed and one disturbed habitat not included in the continental and global studies namely dunes, wetlands (bogs, mires and fens) and greenspaces (green urban areas) (**Supplementary Note Table 2**). In agreement with the continental and global studies of soil prokaryotic diversity we found high alpha-diversity in croplands, but with no significant difference to the one found in grasslands (p=1). However, the urban area harboured a significantly larger alpha diversity than both croplands (p<0.001) and grasslands (p=0.004). Furthermore, no significant difference was observed when comparing wetlands to croplands (p=0.81) or grasslands (p=0.75) (**Supplementary Note Table 2**). Also, in agreement with the continental and global studies we found the lowest alpha-diversity in forests, which in our case was not significantly different to the one observed in dunes (p=0.84) or shrublands (p=1). Conversely, when exploring the investigated terrestrial habitat types, we found the lowest estimated gamma diversity in croplands (highly-disturbed) [N = 93, 95% CI: 18,266-18,487]. The green urban areas (moderately-disturbed) showed significantly higher diversity than the croplands [N=24, 95% CI: 30,158-30,853]. Also, the greenspaces had significantly higher gamma diversity than both forests (less-disturbed) [N=76, 95% CI: 27,819-28,315] and dunes [N=17, 95% CI: 24,387-,25,626], but lower than grasslands (moderately-disturbed) [N = 55, 95% CI: 37,201-37,887] and wetlands (less-disturbed) [N=36, 95% CI: 42,964-,44,027].

**Supplementary Note Table 2. Comparison of alpha-diversity.** Differences in alpha diversity of MFDO1 habitats for Denmark (MFD), as well as published results for Earth (EMP[30]) and Europe (LUCAS[31]). For MFD alpha diversity was measured as observed ASV richness and the statistical analysis conducted with a one-way ANOVA and the TukeyHSD post hoc test ($p<0.05$). For EMP alpha diversity was reported as observed OTU richness and the statistical analysis conducted with a one-way ANOVA and the TukeyHSD post hoc test[30]. For LUCAS alpha diversity was reported as observed zOTU (the same as ASV) richness and the statistical analysis conducted with a one-way ANOVA on ranks (Kruskal-Wallis) and a Wilcoxon post hoc test[31]. Letters (**Sig**) indicate significant differences between groups where groups that share a letter are not significantly different from each other. For the LUCAS study the letters have been reversed and the original reported in brackets.

| **α-diversity** | **MFD** | **Sig** | **EMP[30]** | **Sig** | **LUCAS[31]** | **Sig** |
|---|---|---|---|---|---|---|
| Low ↕ High | Dunes | d | | | | |
| | Forests | d | Forest | bc | Coniferous wood | d (a) |
| | | | | | Broadleaved wood | c (b) |
| | Heath and scrub | cd | Shrubland | abc | | |
| | Grassland formations | b | Grassland | ab | Extensive grassland | b (c) |
| | | | | | Intensive grassland | abc (bcd) |
| | Bogs, mires and fens | bc | | | | |
| | Fields | b | Agriculture | a | Non-permanent crop | ab (cd) |
| | | | | | Permanent crop | a (d) |
| | Greenspaces | a | | | | |

### Supplementary Note 3: Microbes as habitat descriptors

Habitats have historically been described by macroflora and abiotic observations. We investigated how well the microbial community inhabiting those habitats agree with such categories (**Figure 4**). For this analysis we used the 16S rRNA gene fragments from the 10,686 metagenomes classified by the MFG 16S rRNA gene database and used microbial counts as predictor variables for the habitat categories. A total of 25 combinations between predictors, i.e., five taxonomic levels (Phylum n=72, Class n=191, Order n=580, Family n=1493, Genus n=6613), and targets, i.e., five description levels (Sampletype n=3, Areatype n=8, MFDO1, n=16, MFDO2 n=30, MFDO3 n=35), were used to model the data with a random forest algorithm. Overall, the use of higher taxonomic detail yielded better results while the opposite was true for the ontology detail, with a sharp downturn after MFDO1 (**Supplementary Note Figure 1a**). Starting at the highest level of the ontology, i.e., Sampletype, the models are good at discerning between Water, Sediment and Soil (PR AUC of 0.91, 0.91 and 0.99, respectively). However, at the Areatype level the models have different performances on different classes. For instance, the "Urban" soils are harder to recognise (PR AUC=0.60) than both "Natural" and "Agricultural" ones (PR AUC of 0.93 and 0.98). Evidently the classes associated with a niche or specialised microbiome obtain the highest or even perfect scores ("Saltwater" and "Wastewater" in **Figure 4**).

There is information also in the failure of the classification as especially the samples from "Grassland formations", "Greenspaces" and "Fields" are misclassified as each other (**Supplementary Note Figure 1b**). Likewise, the samples from the two freshwater sediment classes ("Urban" and "Natural") are more often misclassified as each other as well as "Bogs, mires and fens" than expected by chance (**Supplementary Note Figure 1b**). There could be several explanations to these misclassifications, such as i) initial erroneous classification in the metadata, ii) spatial or temporal transition between environments. The habitats were provided as a categorical choice, whilst it has been argued that habitats should be considered as a bland of continuous gradients[153]. Similarly, temporal changes of habitats might affect the above-ground flora, used for habitat classification, faster than the below-ground microbiome (which adapts over centuries[154]). Moreover, unless specific techniques are applied, it is impossible to distinguish DNA from live cells and "relic" DNA, possibly making this "lagging microbiome" effect stronger[155].

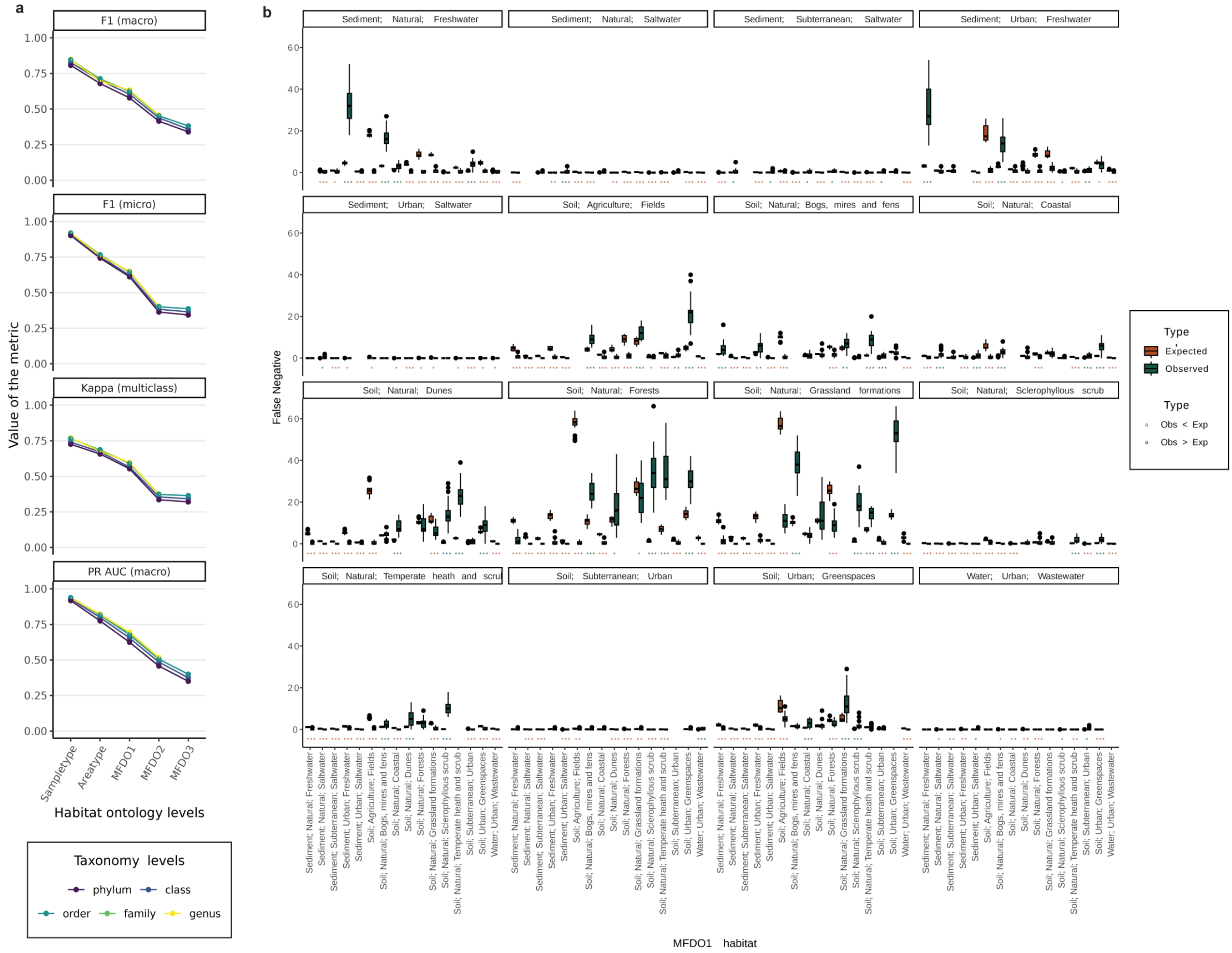


**Supplementary Note Figure 1. Habitat classification metrics. a.** The quality of the classification models was assessed using the F1 (macro and micro), Kappa and the Precision Recall Area Under the Curve (PR AUC) metrics for each model. The points indicated mean values of the metrics per model (n=25: 5 folds x 5 iterations) and the bars indicate the standard deviations. The lines connect measurements from the same taxonomic level, also represented by the colour. According to all the metrics, the performances of the models increased with the increase of taxonomic granularity. The metrics also showed a steady decline in quality along the ontology levels with a particular decrease after MFDO1. **b.** The False Negatives (FN) from the classification models using Order level taxonomy and MFDO1 level ontology labels show that wrong classification is often following habitat-specific patterns. Each facet contains the FN values referring to one of the 16 classes modelled in **Figure 5**. For each wrong assignment (i.e., the classes on the x axis) we computed the expected number of FN given the total number of FN for the reference class in each split and iteration, scaled by the class sizes in each split and iteration. For each "Expected" and "Observed" pair we computed a two-tailed t-test and reported the p-value bracket ("***" for $p < 0.001$, "**" for $0.01 < p < 0.001$ and "*" for $0.05 < p < 0.01$) and coloured according to the sign of the test statistic to indicate which distribution is greater. The samples of several classes are more easily mistaken for some classes than others, e.g., "Soil; Natural; Grassland Formation" is misinterpreted as "Soil; Urban; Greenspaces" significantly more than expected.

### Supplementary Note 4: Few genera make up the majority of the prokaryotic communities

Determining whether a habitat has a characteristic or core microbiome may give insight into the important microbial functions involved in maintaining ecosystem services or adapting to environmental changes[156]. We wanted to determine whether we could determine MFDO1 habitat specific genera, by assessing how prokaryotic taxa were sorted and shared between the MFDO1 habitats. Across Denmark the core population of each MFDO1 habitat ranged from 4 ("Drinking water") to 60 ("Fields") genera (**Supplementary Note Figure 2**). However, only 10 of the 60 genera identified in "Fields" were unique to that habitat type. "Fields" also showed the largest overlap of nine core genera with the other disturbed soil habitat "Greenspaces" including MFD_g_4907 from *Nitrososphaeraceae*, a family of ammonia-oxidising archaea (AOA). The overlap of genera between the disturbed soil environments suggests that the shared ecological space shaped by land management perturbation and cultivation practices (herb-domination, fertilisation and drainage), might have selected for the same genera, including ones associated with nitrogen cycling, different from those present in their natural counterparts. The largest number of habitat-specific core genera were found in "Wastewater", "Biogas", as well as surface and subterranean sediments from "Saltwater". In the case for subterranean saltwater sediments 33 of the 48 core genera had placeholder names and made up 17.8% of the relative abundance (data not shown). Genera with placeholder names are of particular interest as their presence might reflect habitats still underrepresented in the databases but also gives targets for future evaluation.

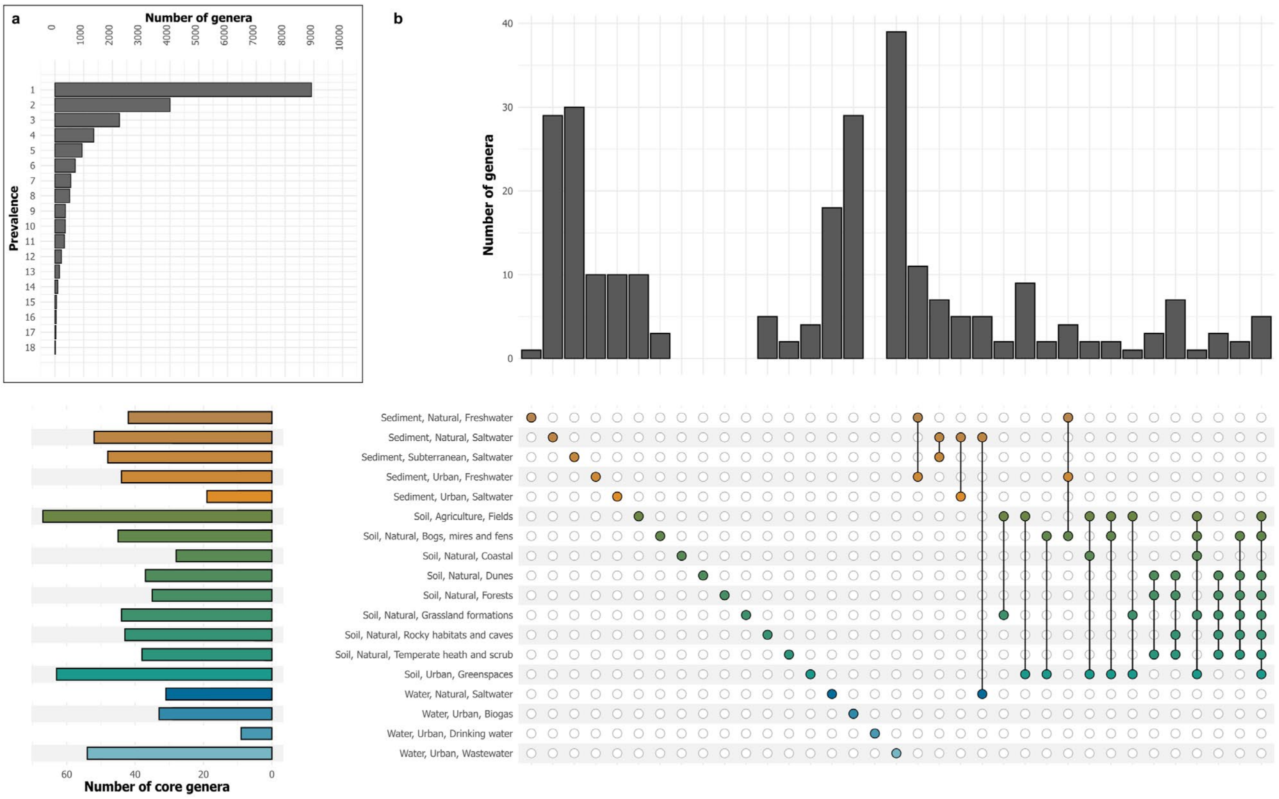


**Supplementary Note Figure 2. Core genera across MFDO1 habitats. a.** The prevalence of all identified genera of the geographically balanced dataset was evaluated by the number of different MFDO1 habitats in which each genera was observed **b.** Upset plot of core genera identified across the 18 MFDO1 habitats.

**Supplementary Note 5: Improving HMM packages for nitrification related genes**

Gene-centric identification of nitrifiers was based on the ammonia monooxygenase subunit A (*amoA*) and nitrite oxidoreductase subunit A (*nxrA*) genes. To improve the resolution and fidelity of this approach, HMM search and classification packages were built or improved. The databases were made using sequences linked to taxonomic information from GTDB Refseq release 214 and the GEM database, and supplemented by genes identified in our MAG database. All genes were translated into protein sequences for protein phylogenetic trees and generation of HMMs. The packages were curated by investigating the protein-phylogeny and taxonomy. In cases where the protein-phylogeny was not in accordance with the expected taxonomic classification of the genome (either isolate or MAG), the genome was evaluated for contamination using DIAMOND blastx[132] v2.0.9.147. This was supplemented by looking at the multiple sequence alignment. Sequences that were partial or appeared to be placed on contaminated contigs, were removed.

The bacterial *amoA* package was based on an existing *amoA/pmoA* package by Singleton et al. 2018[157]. No pre-existing packages were available for archaeal *amoA* or *nxrA/narG.* Within the bacterial *amoA* package, sequences encoding hydrocarbon monooxygenases and PmoA or PxmA from *Methylococcaceae* and *Methylomonadaceae* were clustered at 95% aa identity. The remaining sequences were clustered at 100% aa identity. All archaeal AmoA sequences were clustered at 100% aa identity. Putative AmoA sequences from p__Thermoplasmata; o__RBG-16-68-12 were considered as an outgroup, lacking confirmation of AmoA activity. Remaining AOAs were members of o_Nitrososphaerales, either f_Nitrososphaeraceae or f_Nitrosopumilaceae.

NXRs form NOB and anammox organisms are either facing the cytoplasmic or periplasmic side of the cell membrane, or are localised in the anammoxosome. Cytoplasmic NXRs (cNXR) are homologous to membrane bound cytoplasmic-faced respiratory nitrate reductases (Nar)[158,159]. Denitrification is taxonomically widely distributed[160,161], resulting in more than 10,000 unique (100%aa identity) NarG sequences obtained from GTDB Refseq release 214 and the GEM database. While it is necessary to have representation of NarG for package differentiation between NarG and cNxrA, it is essential to balance the HMM. This was done by clustering NarG sequences from the phyla Bacillota, Actinomycetota, and Firmicutes at 90% aa identity, and NarG sequences from Gammaproteobacteria at 95% aa identity. The remaining NarG/cNxrA sequences were clustered at 100% aa identity, reducing the NxrA/NarG HMM to contain 1390 sequences, of which 1108 are NarG and 282 were cNXR from *Nitrobacter*, *Nitrococcus*, and *Thiocapsa*. Furthermore, *Nitrolancea hollandica*[162] NxrA was included, but clustered with gammaproteobacterial NarG sequences.

The pNXR package consisted of *Nitrospira* (including Palsa-1315), *Nitrotoga*, *Nitrohelix*, *Nitrospina*, along with members of the Nitrospirales order, f__NS-4; g__NS-12, and f__UBA8639; g__UBA8639. Anaerobic ammonia oxidation (anammox) pNXR sequences were also present in the pNXR package. However, no Anammox (*Anammoxibacter*, *Scalindua*, and *Brocadia*) were detected with single copy marker genes, except in low abundance in 'Wastewater' (**Supplementary Note Figure 3**). Reads from the shallow metagenomes did, however, attract to the anammox clades containing "NXR/NAR" sequences from GTDB and GEM databases. These reads were investigated using blastx against nr (Jan 2024), and showed high sequence identity to proteins from non-anammox bacterial genera, probably due to the homology between bacterial NarG and anammox NXR[158]. On this basis, gene-centric identification of anammox remains unresolved within this study. All sequences in the pNXR package were clustered at 100% sequence identity, resulting in a package-size of 192 sequences.

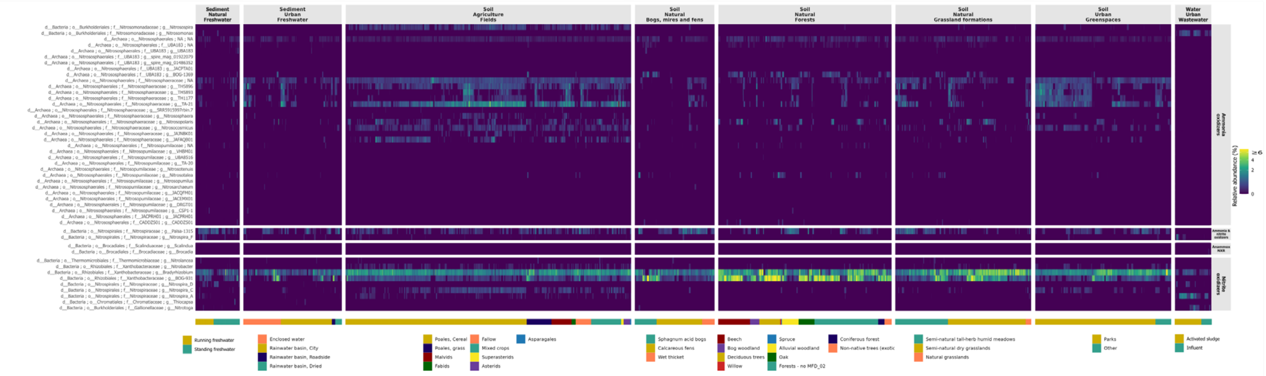

**Supplementary Note Figure 3. Nitrifier distribution in Danish habitats based on SingleM. a.** The Principal Coordinate Analysis (PCoA) of nitrifiers based on Hellinger-transformed Bray–Curtis Distance on the relative abundances of nitrifiers, faceted into 8 selected MFDO1 habitats. Habitat colour codes are the same as panel b. **b.** Heatmap showing the distribution of nitrifiers across 8 selected MFDO1 habitats in MFD short-read metagenomes based on SingleM with metapackage supplemented with short-read metagenome recovered MAGs.

Screening the shallow metagenomes showed high attraction of reads to *Nitrososphaeraceae* AmoA (**Supplementary Note Figure 4**). AOAs within *Nitrososphaeraceae* have been observed in high relative abundance in soil, but are poorly understood[163,164]. AmoA-like sequences from *Nitrososphaeraceae* MAGs were filtered based on length > 200 aa, and de-replicated at 100% aa identity. Adding these sequences allowed for the generation of two separate clades for g_TH5893, in which only a single AmoA had been identified in publicly available genomes (GEMOTU_03734). With the additions from MFD, it was possible to identify g_TH5893 across agricultural soils, grasslands and greenspaces (**Supplementary Note Figure 4**). Furthermore, an additional 10 sequences from g_TH5896 could be added to the GraftM package, allowing for generation of two separate clades, TH5896_1 and TH5896_2 (**Supplementary Note Figure 4, Supplementary Note Figure 5**). Only two partial AmoA sequences were present in g_TH5896 prior to the addition of MFD sequences. The sequences added to clade 1 allowed for improved resolution and identification of the potential AOA in Danish soil and sediment of various habitat types, especially field samples

(**Supplementary Note Figure 4**). To assess the attraction of reads to the novel clades of AOA AmoA, we evaluated the coverage profile across the HMM, which showed an even distribution across the 216 aa (**Supplementary Note Figure 5**). It was evident from both protein phylogenetic placement trees and coverage that the majority of reads are assigned to the *Nitrososphaeraceae* umbrella group and the TA21/TA21_*Nitrosopolaris* cluster (**Supplementary Note Figure 5**).

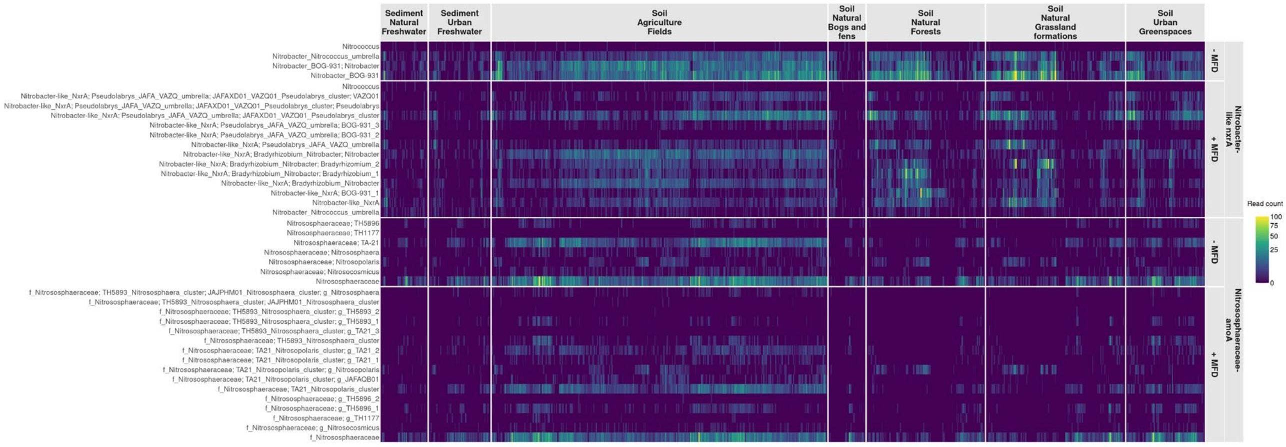


**Supplementary Note Figure 4. The effect of GraftM package update on nitrifier habitat distribution.** Heatmap showing the reads per kilobase million (RPKM) of nitrification genes, faceted by MFDO1. Samples are sorted by hierarchical clustering. Assignment of reads to each group before (-MFD) and after (+MFD) additions of AmoA or NxrA sequences from SR-MAGs obtained in this study.

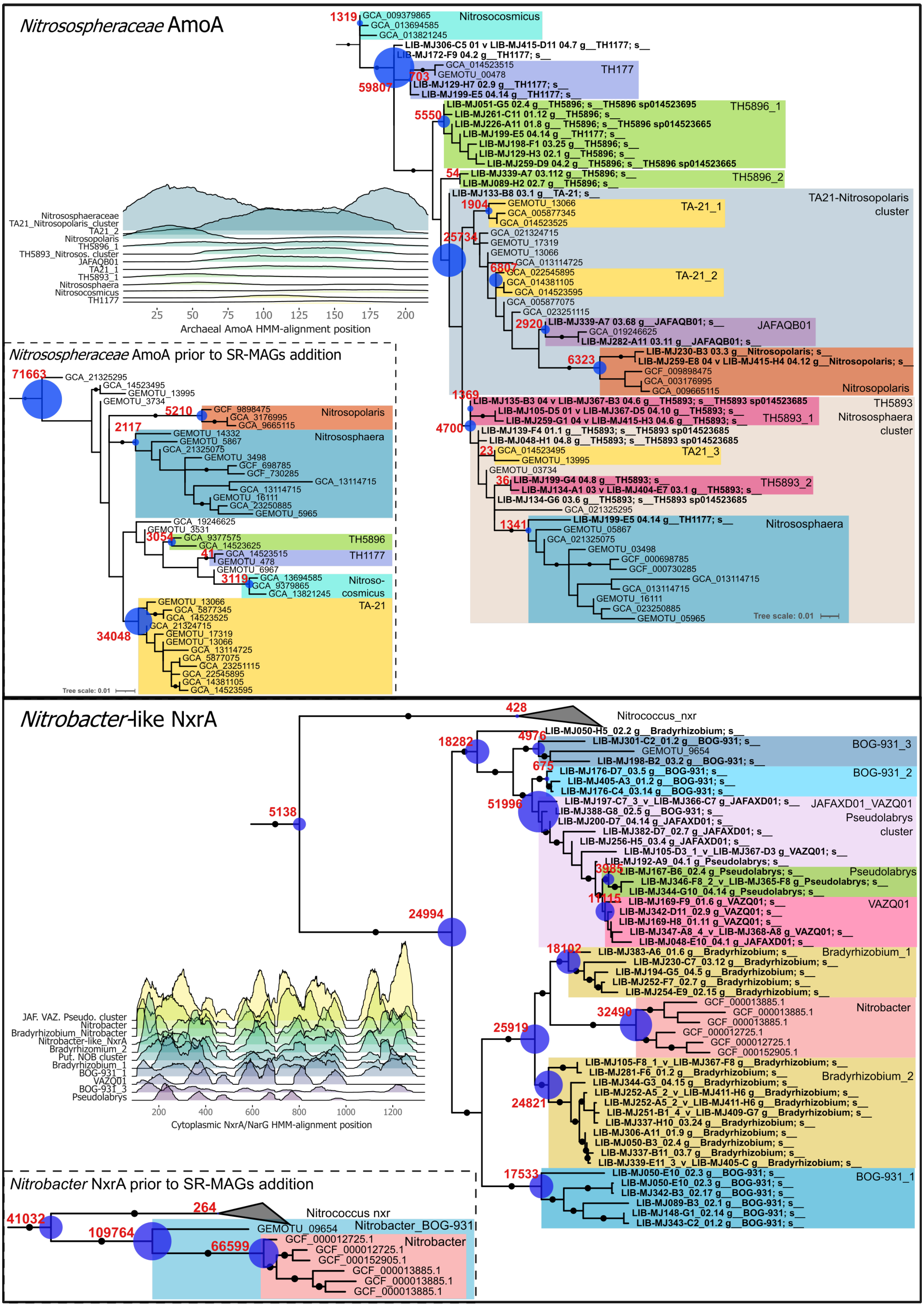

Nitrososphaeraceae AmoA
Nitrosocosmicus
TH177
TH5896_1
TH5896_2
TA21-Nitrosopolaris cluster
TA-21_1
TA-21_2
JAFAQB01
Nitrosopolaris
TH5893 Nitrososphaera cluster
TH5893_1
TA21_3
TH5893_2
Nitrososphaera
Archaeal AmoA HMM-alignment position
Nitrososphaeraceae AmoA prior to SR-MAGs addition
TH5896
TH1177
Nitroso-cosmicus
TA-21
Tree scale: 0.01
Nitrobacter-like NxrA
Nitrococcus_nxr
BOG-931_3
BOG-931_2
JAFAXD01_VAZQ01 Pseudolabrys cluster
Pseudolabrys
VAZQ01
Bradyrhizobium_1
Nitrobacter
Bradyrhizobium_2
BOG-931_1
Cytoplasmic NxrA/NarG HMM-alignment position
Nitrobacter NxrA prior to SR-MAGs addition
Nitrococcus nxr
Nitrobacter_BOG-931

**Supplementary Note Figure 5. Attraction of reads to updated nitrifier protein-phylogenetic trees.** Protein phylogeny of the *Nitrosospheraceae* family of AOAs (top), and *Nitrobacter*-like NxrA/NarG sequences (bottom). In red are the number of reads from the all shallow metagenomes assigned to each node identified with GraftM. The size of the blue circle is scaled to the relative number of reads. The genome from which the AmoA or NxrA/NarG sequence was inferred is indicated at the leaf of the tree. In bold are AmoA or NxrA/NarG sequences from SR-MAGs obtained within this study, supplemented with the GTDB-Tk classification. The protein-phylogenetic tree before addition of AmoA or NxrA/NarG sequences from SR-MAGs is present in the lower left corner for each panel. Coverage profiles are displayed for the clades depicted in the tree to visualise the alignment of reads along the HMM.

A large proportion of nitrification genes from the shallow metagenomes also attracted to the *Nitrobacter* NxrA clade (**Supplementary Note Figure 4**). *Nitrobacter*-like NxrA sequences were extracted from MAGs, filtered based on length >600 aa, and dereplicated against the NxrA/NarG sequences within the GraftM package. Sequences obtained from MAGs classified as unknown species within the genera *Pseudolabrys*, JAFAXD01, and VAZQ61 were clustered at 99% aa identity, while those from BOG-931 and *Bradyrhizobium* were clustered at 100% aa identity. Both BOG-931 and *Bradyrhizobium* formed several clades within a protein phylogenetic tree (**Supplementary Note Figure 5**)**.** The NxrA-like clade BOG-931_1 formed a monophyletic clade in close proximity to *Nitrobacter* and *Nitrococcus* NxrA, and had a strong signal relating to habitat types (**Supplementary Note 6**, **Supplementary Note Figure 4**)**.** Clade BOG-931_3 showed similar patterns as BOG-931_1, but attracted fewer reads and formed a less strict group in the phylogenomic tree. While sequences assigned to *Nitrobacter* still appeared in fields and greenspaces, it was evident that sequences previously assigned to *Nitrobacter*, could now be ascribed to clades of *Nitrobacter*-like NxrA sequences within undescribed genera of the *Xanthobacteriaceae* family (**Supplementary Note Figure 4**). This was prominent for clusters of NxrA-like sequences within *Pseudolabrys*, JAFAXD01, and VAZQ61, and clades of *Bradyrhizobium* in field samples. NxrA-like sequences from the genus BOG-931 could be identified in natural soils, in particular forests and grasslands. Using single copy marker genes to determine relative abundance showed nearly no *Nitrobacter* present in Danish soils (**Figure 6**). Whether the sequences still assigned to *Nitrobacter* NxrA do originate from *Nitrobacter* remains to be determined. We also investigated the coverage profile of reads assigned to the HMM (**Supplementary Note Figure 5**). Reads assigned to *Nitrobacter*-like NxrA were not mapping to the entire HMM. The normalisation to RPKM takes the length of the HMM into account (1350 aa), reducing the RPKM relative to other nitrifier groups where sequences align to a broader range of the HMM, and should be taken into account when comparing read abundance across groups. By including sequences from MAGs, we see a markedly improved resolution of the cNXR, as we are now able to differentiate highly abundant sequences formerly ascribed to *Nitrobacter* (**Supplementary Note Figure 5**).

The MAGs were also screened for potential additions to the *Nitrospira* pNxrA/NarG HMM, since *Nitrospira*-like nxrA sequences assigned to *Nitrospira* and Palsa-1315 umbrella clades accounted for a significant proportion of pNXR sequences (**Figure 6d**). This was also reflected in the discrepancies observed between identification with nitrifier genes and single-copy marker genes (**Figure 6d and 6e**). A total of nine NxrA sequences originating from novel species within Nitrospira_F, Nitrospira_D, Nitrospira_A, and Palsa-1315 could be added (data not shown). Topology of the tree changed only slightly upon the addition of *Nitrospira* NxrA sequences. While resolution of CMX *Nitrospira* clade B (Palsa-1315) was improved markedly using our search database, the resolution of canonical *Nitrospira* and CMX *Nitrospira* clade A (Nitrospira_F) remained unresolved, and could not differentiate the other genera belonging to the former *Nitrospira* group (Nitrospira A to F).

We also examined the presence of known habitat specific nitrifiers to evaluate the HMM models. This includes the AOA *Nitrosotalea,* which was detected in the acidic bogs (**Figure 6d**), inline with their characterisation as obligate acidophiles[165–167]. Urban wastewater showed prevalence of the expected AOB *Nitrosomonas*, and the often associated NOB *Nitrospira_A* (encompassing *N. defluvii*) in influent and activated sludge (**Figure 6d**). As expected, *Ca*. Nitrotoga was detected in the activated sludge. Furthermore, wastewater had clustering distinct from the other habitats (**Figure 6a**).

### Supplementary Note 6: Metabolic potential of novel nitrifiers

NxrA sequences assigned to *Nitrobacter* were identified across nearly all agricultural soil samples, and were also present in most forest, grassland and greenspace soils (**Supplementary Note Figure 4**). Prior to HMM package improvement with sequences from the MAGs, *Nitrobacter nxrA*-like genes were classified as *Nitrobacter*, providing an inaccurate representation of *Nitrobacter's* abundance and distribution. The sequences actually belong to other members of the *Xanthobacteraceae*, primarily *Bradyrhizobium* spp., and the uncharacterised genus BOG-931. We examined the topology of the protein trees in comparison to the topology of the MAGs, and identified that distinct clades of multiple species within these genera encoded the *Nitrobacter*-like NxrA (**Supplementary Note Figure 6**). In contrast, no MAGs containing *Nitrobacter*-like NxrAs were assigned to the *Nitrobacter* genus. In particular, a monophyletic clade of *Nitrobacter*-like *nxrA* from BOG-931 grouped close to *Nitrobacter* and *Nitrococcus nxrA*, while the associated genomes also clustered together in a phylogenomic tree (**Supplementary Note Figure 6**).

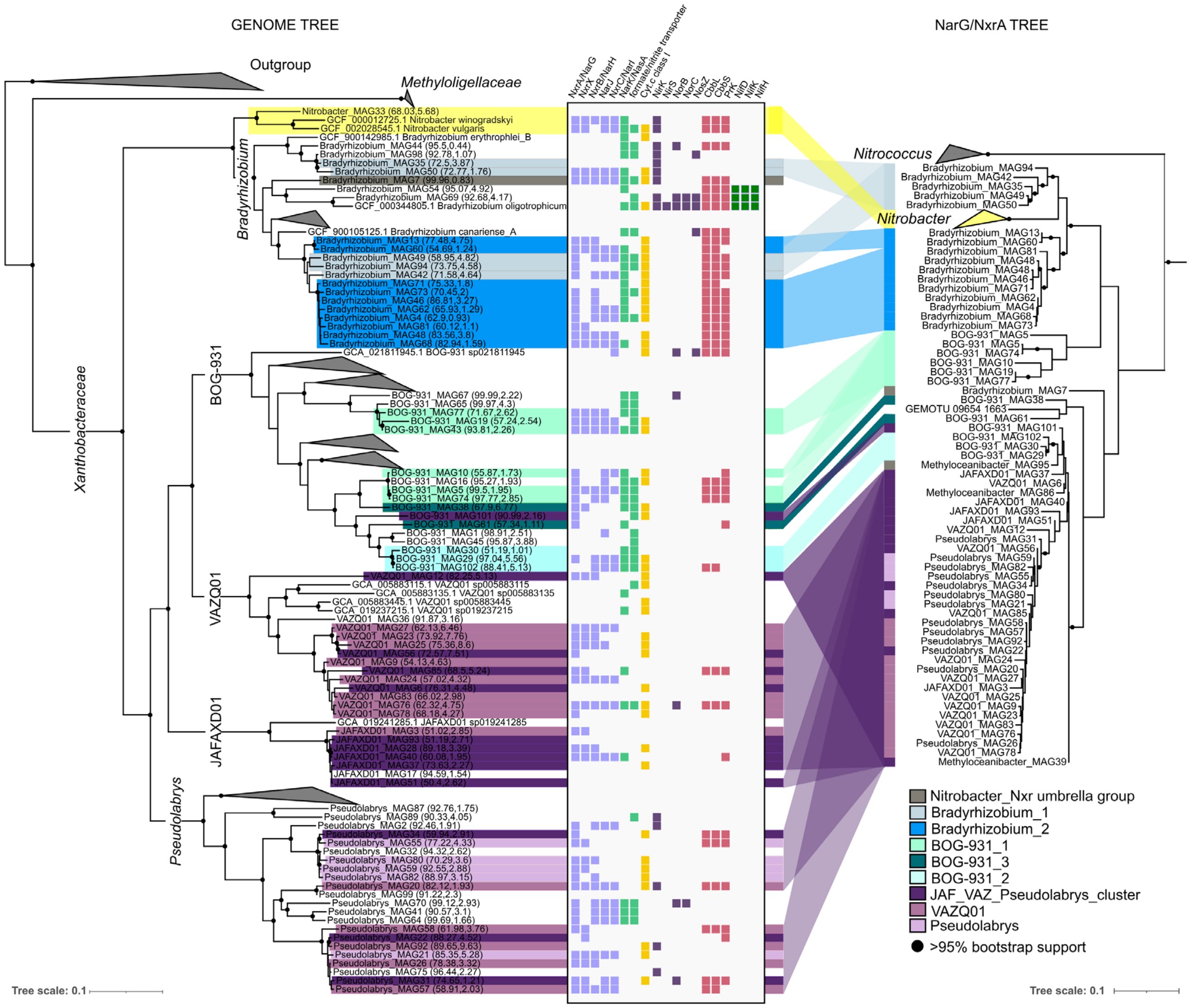


**Supplementary Note Figure 6. Aligned genome and protein tree of new *Nitrobacter*-like *narG/nxrA* encoding MAGs.** Genome tree (left) of potential nitrite oxidisers containing *Nitrobacter*-like NxrA, belonging to the *Xanthobacteriaceae*, primarily *Bradyrhizobium* spp., genus BOG-931 spp., and *Pseudolabrys* spp. Protein tree (right) of NxrA sequences from *Nitrococcus*, *Nitrobacter*, and *Nitrobacter*-like NxrA recovered from SR-MAGs. Wedges represent clusters of genomes within the genera that do not contain the NxrA-like sequences of interest. Protein subunits involved in nitrite oxidation or reduction are indicated by: NxrA/NarG, NxrX, NxrB/NarH, NarJ, NxrC/NarI, nitrite/nitrate transport: NarK/NasA, formate_nitrite_transporter, electron transfer: Cyt.c class I, nitrite reduction: NirK, NirS, nitric oxide reduction: NorB, NorC, nitrous oxide reduction: NosZ, carbon dioxide fixation: CbbL, CbbS, PrK, nitrogen fixation: NifD, NifK, NifH.

This clade was notably more abundant in forests and grasslands compared to other habitat types (**Supplementary Note Figure 4**). We investigated the MAGs for autotrophy signal similar to *Nitrobacter*, specifically C fixation using the CBB cycle, and found CBB cycle genes (form 1 ribulose bisphosphate carboxylase/oxygenase, large and small subunits determined by NCBI BLAST against nr (Feb 2024)) in the *Bradyrhizobium*, which is expected for this genus of functionally diverse facultative autotrophs[168]. Interestingly, the clade of BOG-931 encoding the putative NXR also encoded CBB cycle genes (also form 1), which was uncommon for the MAGs belonging to this genus in general (**Supplementary Note Figure 6**, **Supplementary Note Figure 7**). The *Nitrobacter* and *Nitrococcus* cytoplasmically oriented NXR is a nitrate reductase that works in reverse[169]. Since novel NXR sequences could represent either new NOB or new nitrate reducers, we investigated the gene synteny and possible evolutionary context by including contiguous long-read (LR) assembled HQ MAGs of BOG-931 (unpublished).

Of 131 LR-HQ BOG-931 MAGs, only 4 encoded a *Nitrobacter*-like *nxrA* (**Supplementary Note Figure 7**). Gene synteny of LR-HQ MAGs of BOG-931 resembled the *nxr/nar* operon of *N. winogradskyi* and *N. hamburgensis*[170,171], consisting of cytochrome c class I, *nxrA*, *nxrX*, *nxrB/narH*, *narJ*, *narI* (**Supplementary Note Figure 7**). Cytochrome c oxidase gene clusters were found in all genomes, and were located in close proximity to *nxr/nar* operon (less than 5 kbp) in 2/4 MAGs. However, the copper-containing nitrite reductase (*nirK*) found in *Nitrobacter*[171] was not identified in the BOG-931 MAGs. The *nxr/nar* operons were flanked by transposases in all genomes (**Supplementary Note Figure 7**), and could suggest the acquisition of the *nxr/nar* operon by horizontal gene transfer. All MAGs containing *Nitrobacter*-like *nxrA* also encoded at least one copy of a formate/nitrite transporter. CBB-cycle indicators RuBisCo type I large and small subunits (*cbbL, cbbS*) were encoded in close proximity to the *nxr/nar* operon in one MAG (~5 kbp), along with a NarK-like nitrate/nitrite transporter (**Supplementary Note Figure 7**). Furthermore, *cbbL, cbbS* were located 1.4 Mbp from the *nxr/nar* operon in a circular BOG-931 MAG. RuBisCO operons were identified in 2/4 HQ LR-MAGs, and in 4/5 SR HQ MAGs that encode *Nitrobacter*-like *nxrA*. In contrast, none (0/127) of the HQ LR BOG-931 MAGs encoded RuBisCO without *Nitrobacter*-like *nxrA* (**Supplementary Note Figure 7**). While the assimilation of carbon through the CBB-cycle is conserved in *Nitrobacter*, RuBisCO have been reported to be localised on plasmids in *N. hamburgensis*[171]. Based on gene phylogeny and operon structure, the RuBisCO encoded in BOG-931 MAGs appeared to be the IC form, being flanked by *cbbX* and other *cbb* metabolic genes, and lacked carboxysome genes, similar to what is observed in *Nitrosococcus oceani*, *Nitrosospira multiformis* and *Bradyrhizobium japonicum*[172].

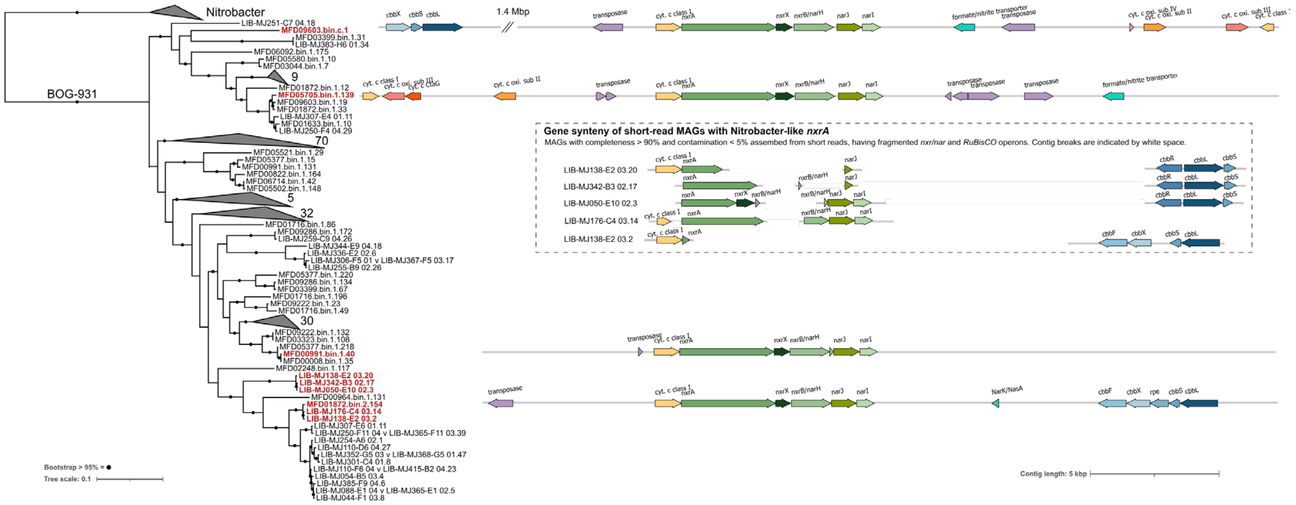


**Supplementary Note Figure 7. Genome tree of BOG-931 SR and LR MAGs of high quality.** Genome tree (left) of potential nitrite oxidisers containing *Nitrobacter*-like *narG/nxrA*, belonging to the genus BOG-93. LR-MAGs (MFDxxxxx.bin.xx) included are of HQ based on the MIMAG standard, while SR-MAGs (LIB-MJxxx-xx) with completeness >90% and contamination <5% have been included. Genomes in red encode *Nitrobacter*-like *nxrA* sequences. For LR-MAGs, gene synteny is only displayed for contigs that also encode Nitrobacter-like *nxrA*. In the circular genome MFD09603.bin.c.1, RuBisCO is present on the same contig, with a distance between *nxr/nar* and *RuBisCO* operons of 1.4 Mpb. Other nitrification genes on the same contig as Nitrobacter-like NarG/cNXR are displayed. In the box (dotted line) are *nxr/nar* genes from SR-MAGs, along with *RuBisCO*. Here, genes are located on several contigs. The arrangement of genes from different contigs is done manually for visual comparability. Wedges represent clusters of genomes within the genera that do not contain the NxrA-like sequences of interest, displaying the number of genomes collapsed into each clade.

Prior to our MAG recovery, only 4 species for BOG-931 were present in GTDB (v214), and we have added an additional 69 species. Confirmation of BOG-931, and other members of the *Xanthobacteraceae* encoding *Nitrobacter*-like *nxrA,* as potential NOB or incomplete denitrifiers would require culturing to determine if the identified NAR works in reverse as an NXR, rather than as a typical nitrate reductase. Nevertheless, identification and distinction of these putative groups from *Nitrobacter* is an important step towards improved exploration of the nitrogen cycle and the evolution or distribution of NAR and NXR.
